# Enzyme-powered DNA origami nanostructures for enhanced mucosal diffusion

**DOI:** 10.1101/2025.08.29.672641

**Authors:** Matteo Tollemeto, Emily Tsang, Lars J.M.M. Paffen, Lasse H. E. Thamdrup, Jan van Hest, Tania Patiño Padial, Kurt V. Gothelf, Anja Boisen

**Affiliations:** The Danish National Research Foundation and Villum Foundation’s Center IDUN, Department of Health Technology, Technical University of Denmark, Kgs. Lyngby, Denmark; Department of Chemistry and Interdisciplinary Nanoscience Center (iNANO), Aarhus University, Aarhus C, 8000 Denmark; Department of Biomedical Engineering, Institute for Complex Molecular Systems, Eindhoven University of Technology, Eindhoven, The Netherlands

**Author notes:** **Corresponding authors:** Matteo Tollemeto, Tania Patiño Padial, Kurt V. Gothelf. These authors contributed equally to the work (Matteo Tollemeto, Emily Tsang).

**Keywords:** Single-particle tracking, Oral drug delivery, Nanomotors, FRET

## Abstract

Crossing mucosal barriers is a central challenge for oral drug delivery, where nanoparticle design must balance stability with mobility in complex fluids. Here, we demonstrate DNA origami as a programmable platform to investigate these processes. Using FRET analysis, we show that DNA nanostructures retain their structural integrity for extended periods in porcine intestinal fluid and mucus, establishing their suitability for biologically relevant environments. Building on this, we used single-particle tracking to assess enzyme-powered propulsion within mucus. Both urease and catalase enhanced diffusion only when anchored to the DNA origami structure, with propulsion persisting for tens of minutes. Importantly, enzyme spatial organization dictated performance: symmetric urease placement improved mobility via uniform local pH shifts, while asymmetric catalase placement enabled efficient bubble-driven propulsion. These results highlight DNA origami as a uniquely versatile tool to dissect structure-function relationships in mucus transport and provide design principles for next-generation, enzyme-powered oral delivery systems.

## 1. Introduction

Nanomotors are increasingly recognized as promising tools for overcoming biological barriers in drug delivery.^1–4^ Early catalytic nanomotors often relied on reactive metals such as zinc and magnesium, which generate gas bubbles upon contact with aqueous or acidic environments, as well as platinum, which decomposes hydrogen peroxide into oxygen bubbles to propel the particles.^5–7^ While these metals are intrinsically biocompatible, their reliance on high concentrations of fuel limits their applicability.^4,8^ In contrast, the development of enzyme-powered nanomotors, offers improved catalytic efficiency, requires lower fuel concentrations, and expands the range of biocompatible substrates beyond peroxide, thereby providing greater control and compatibility within physiological environments. “Enzymes such as urease, catalase, and lipase enable self-propulsion via chemical gradients (diffusiophoresis), local modifications of the surrounding environment, or oxygen bubbles, with the dominant mechanism depending on fuel concentration, offering more biologically compatible and controllable propulsion compared to traditional metal-based nanomotors.^4,8–11^

One area where such systems could be transformative is mucosal drug delivery, including applications in the gastrointestinal (GI), respiratory, and reproductive tracts.^8,9,12,13^ These mucosal surfaces are protected by a dense and adhesive mucus layer that traps foreign particles and impedes the penetration of drug carriers.^14–16^ While this barrier plays a vital role in host defense, it also represents a major challenge for effective therapeutic delivery.^17^ In particular, oral drug delivery has long been a preferred route due to the high surface area and absorptive capacity of the small intestine.^18^ However, achieving efficient delivery remains challenging, as the GI mucus also undergoes continuous turnover and contains a variety of digestive enzymes.^15,19^ These features limit the ability of drug carriers to reach and penetrate the epithelial surface. As a result, nanoparticles that can maintain structural integrity and effectively navigate through mucus are of growing interest across multiple mucosal delivery applications.^16,20^

Among the various nanocarrier platforms, DNA origami nanostructures offer unique advantages due to their precise control over size, shape, and surface functionalization. ^21–24^ Recent work has highlighted the potential of DNA origami for use in physiological settings, including our own studies on oral drug delivery, by demonstrating that these structures can be engineered to effectively diffuse through mucus barriers.^25–27^ “Recently, DNA nanostructures have been combined with enzymes to achieve self-propulsion.^28^ However, their application as actively propelled carriers to traverse biological barriers, such as the mucosal layer, remains largely unexplored.

In this study, we first evaluated the stability of DNA origami nanostructures in complex biological fluids before investigating their use as enzyme-powered nanomotors to penetrate the mucus barrier. We focused on two propulsion strategies: urease- and catalase-driven motion within a GI-relevant mucus environment. Urease catalyzes the hydrolysis of urea, releasing ammonia and raising the pH, which triggers a gel-to-sol transition in mucus, reducing viscosity and facilitating transport.^8^ Catalase decomposes hydrogen peroxide into water and oxygen, generating bubbles that physically propel the nanostructures forward.^9^ These enzymes were selected not only for their well-characterized propulsion mechanisms but also because their substrates are upregulated in inflammatory GI conditions, serving as endogenous fuels. Urea levels increase in conditions such as uremia and microbial dysbiosis, while hydrogen peroxide is elevated during oxidative stress in inflamed tissues.^29–31^ This pathophysiological relevance supports the use of urease and catalase as bio-integrated propulsion systems for targeted delivery in inflamed mucosal sites.

While molecular asymmetry has been explored in nanomotor design, particularly by Sánchez et al., it is often overlooked in biologically relevant settings, partly because it is challenging to achieve in a controlled and reproducible manner.^9,32^ Building on this concept, we have already performed work on the programmable positioning of urease enzymes on DNA-based nanostructures, systematically varying the degree of anisotropy. Here, we further exploit the programmability of DNA origami to investigate, for the first time, how surface-bound enzymes influence particle mobility in mucus. By arranging enzymes in symmetric or asymmetric configurations, we examine how different design parameters affect propulsion and transport through this biological barrier.^25^

Together, this work advances the understanding of how DNA origami can be engineered for active transport across mucus barriers, combining insights into structural stability, enzyme-based propulsion, and nanoscale symmetry. By addressing these factors in an integrated manner, our study lays the foundation for developing smart, modular systems for oral drug delivery.

## 2. Results and Discussion

### 2.1. Design and structural integrity of DNA origami in the gastrointestinal tract

One key factor influencing nanoparticle diffusion through mucus is size, with particles smaller than 200 nm generally demonstrating superior mobility across the mucin mesh.^15,17^ To meet this criterion, we employed the M13mp18 scaffold to assemble flat sheet (FS) DNA origami structures with a theoretical size of 102 × 72 nm (height × length). Assembly was achieved by thermal annealing of the scaffold with a library of short, sequence-specific staple strands that direct the folding via complementary base pairing.^33^ Atomic force microscopy (AFM) was used to confirm the successful formation and morphology of the nanostructures. Analysis of 50 individual DNA origami confirmed an average size of 110.1 ± 3.9 nm × 74.4 ± 5.3 nm (height × length), consistent with the intended FS geometry and well within the optimal size range for mucus penetration (**Fig. 1A**).

**Figure 1:**
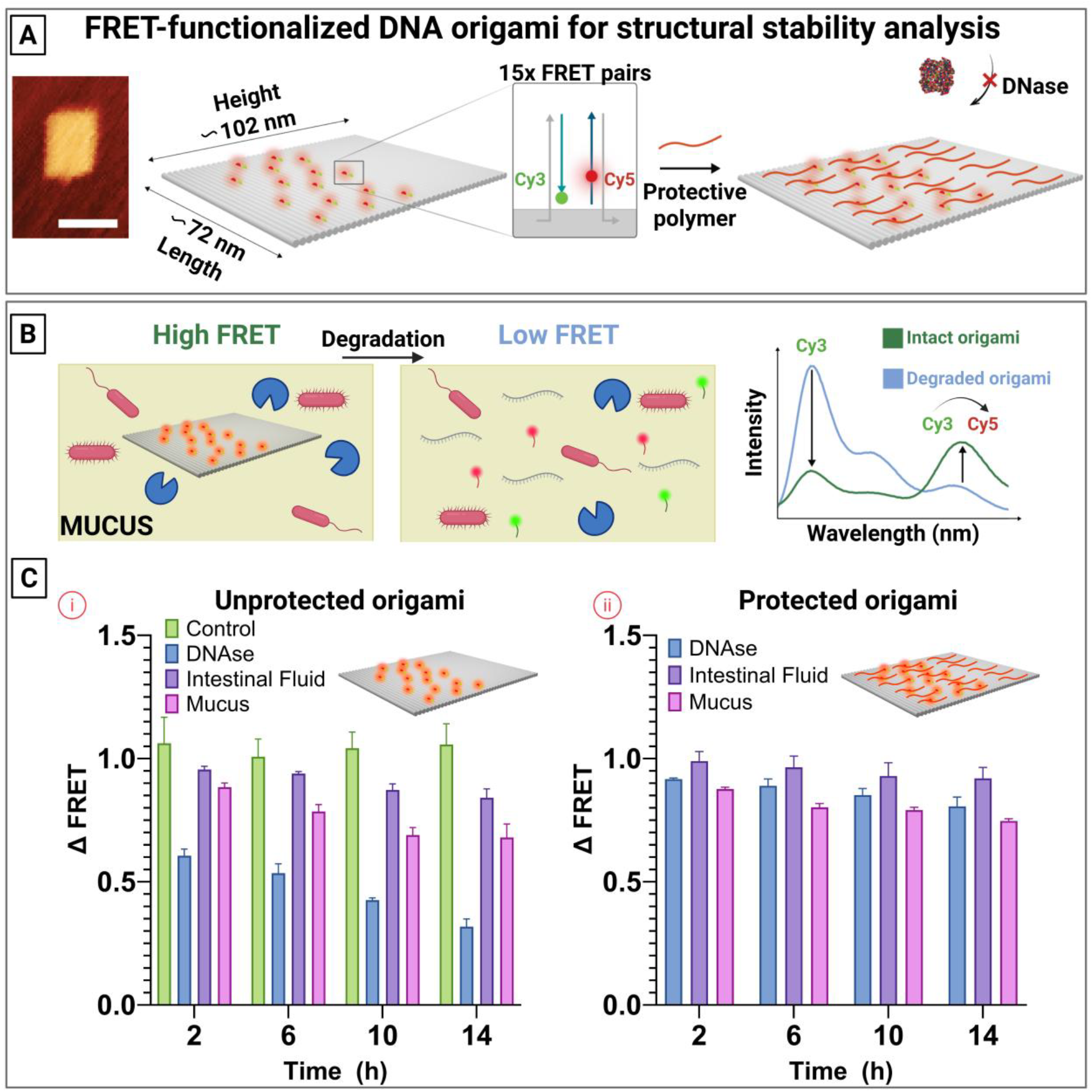
FRET analysis of DNA origami stability in ex vivo intestinal fluids. (A) Schematic illustrating DNA origami assembly with FRET dye pairs and the protective polymer coating designed to prevent degradation (scale bar: 100 nm). (B) Illustration of DNA origami degradation in mucus, showing expected changes in Cy3 and Cy5 emission intensities. (C) (i) Normalized FRET intensity changes over time for uncoated DNA origami, relative to time 0; (ii) corresponding FRET intensity changes for polymer-coated DNA origami, normalized to time 0. Data shown as mean ± SEM (N=4).

While DNA origami structures have recently emerged as promising candidates for oral drug delivery due to their apparent long-term stability, their behavior in biologically relevant environments such as intestinal fluid and mucus remains largely unquantified.^25^ To address this, we employed Förster Resonance Energy Transfer (FRET) to monitor the degradation of various DNA origami designs.^27^ Briefly, each FRET pair consists of two DNA strands, labeled with either Cy3 or Cy5 dyes, which are hybridized to adjacent staple strand extensions spaced approximately 2 nm apart (**Fig. 1A**). As the DNA origami degrades, the distance between the fluorophores increases, leading to a progressive loss of FRET signal.^34,35^ The FRET efficiency was recorded over a period of up to 14 hours and normalized to the initial signal at time zero. This technique, which we have previously validated for assessing structural integrity over time, was re-adapted to assess stability in *ex vivo* porcine intestinal fluid, mucus, as well as high-salt buffer (as a positive control), and in the presence of DNase (as a negative control) (**Fig. 1B**).^27^

The results demonstrate that the control samples in high salts remained stable throughout the 14-hour period, while the DNase-treated negative control showed a rapid decrease in FRET signal, dropping to approximately 60 ± 4% within the first 2 hours and reaching around 32 ± 5% by 14 hours, consistent with our previous findings. Notably, the DNA origami structures incubated in *ex vivo* intestinal fluid and mucus sustained stability over the entire time course. This is likely due to the high ionic strength of both media, which helps preserve the structural integrity of the DNA nanostructures.^36–38^ Mucus and intestinal fluids typically exhibit ionic strengths in the range of 50–200 mM, primarily due to the presence of monovalent ions such as sodium, potassium, chloride, and bicarbonate.^37,38^ Although magnesium, commonly considered the primary stabilizer of DNA origami, is not abundant in these environments, the electrostatic shielding provided by these other salts might contribute to structural stabilization.^39^ Moreover, recent studies have shown that DNA origami can remain stable even at low magnesium concentrations or in the presence of alternative ionic species, supporting their compatibility with physiologically relevant fluids.^40,41^ Quantitatively, approximately 84 ± 6% of the structures remained intact in intestinal fluid and around 68 ± 9% in mucus after 14 hours, indicating that these particles maintain their stability under physiologically relevant conditions (**Fig. 1C i**). These findings suggest that DNA origami may be suitable carriers for therapeutic delivery within the intestinal lumen, in the mucus layer, and potentially even for transport across the mucus barrier. Given that mucosal penetration and epithelial uptake typically occur within a few hours after oral administration, the observed structural stability over 14 hours supports their potential for such applications.^19^

We next incubated the DNA origami with poly(cystaminebisacrylamide-1,6-diaminohexane) (PCD), a cationic polymer previously reported to protect DNA nanostructures in biological environments.^27^ PCD provides a biocompatible and reversible coating that protects the origami during extracellular transit while enabling functional recovery upon cellular uptake. Since the cellular uptake of DNA origami depends on multiple factors, including shape, size, cell type, and the presence of targeting ligands, we sought to test PCD both as a means of protecting the structures from degradation and as a potential strategy for facilitating intracellular delivery in the future.^42^ Once internalized, the polymer is cleaved in the reductive intracellular environment, allowing the DNA origami to regain its native structure and activity.^27^ The resulting polyplexes were formed at an N/P ratio of 1 based on prior findings demonstrating optimal stabilization against DNase-mediated degradation. Consistent with those results, our data showed that PCD-coated DNA origami retained over 80% of their structural integrity in the presence of DNase after 14 hours, compared to only 30% for unprotected structures. Enhanced protection was also observed in *ex vivo* intestinal fluid and mucus, where the stabilized origami showed 91 ± 7% and 75 ± 1% stability, respectively (**Fig. 1C ii**). Overall, our findings indicate that DNA origami structures exhibit notable stability in GI environments such as mucus and intestinal fluid. However, applying protective coatings like PCD can further enhance their long-term stability, offering an additional strategy to improve their performance for oral drug delivery applications.

### 2.2. Analyzing mucosal diffusion of DNA origami via single particle tracking

To evaluate how surface coating affects the diffusion of DNA origami through mucus, we compared the mobility of two distinct designs, the FS and the 14-helix bundle (14HB), in *ex vivo* porcine intestinal mucus using single particle tracking (SPT) (**Fig. 2**).^27,35,43^ We selected the FS (2D - rectangles) and 14HB (rods) for their anisotropic geometries, which allow ligands to be organized in specific regions of the nanostructure, enabling controlled spatial presentation of various ligands and potentially influencing their interaction with mucus.^44^ Experiments were conducted with an Oxford NanoImager (ONI) operating in highly inclined and laminated optical sheet (HiLo) mode. For visualization, each DNA origami shape was labeled with six ATTO 647N fluorophores to enable precise localization. Tracking was performed by recording 10-second videos at 37 °C and 100 frames per second (FPS), allowing us to capture the real-time motion of individual particles (**Fig. 2A i**). We have selected representative particle trajectories (**Fig. 2A ii**), with each trajectory color-coded based on its diffusion coefficient (Deff), ranging from 0 to 1 µm^2^/s. This qualitative mapping gives a visual impression of mobility differences between uncoated and coated FS structures.

**Figure 2:**
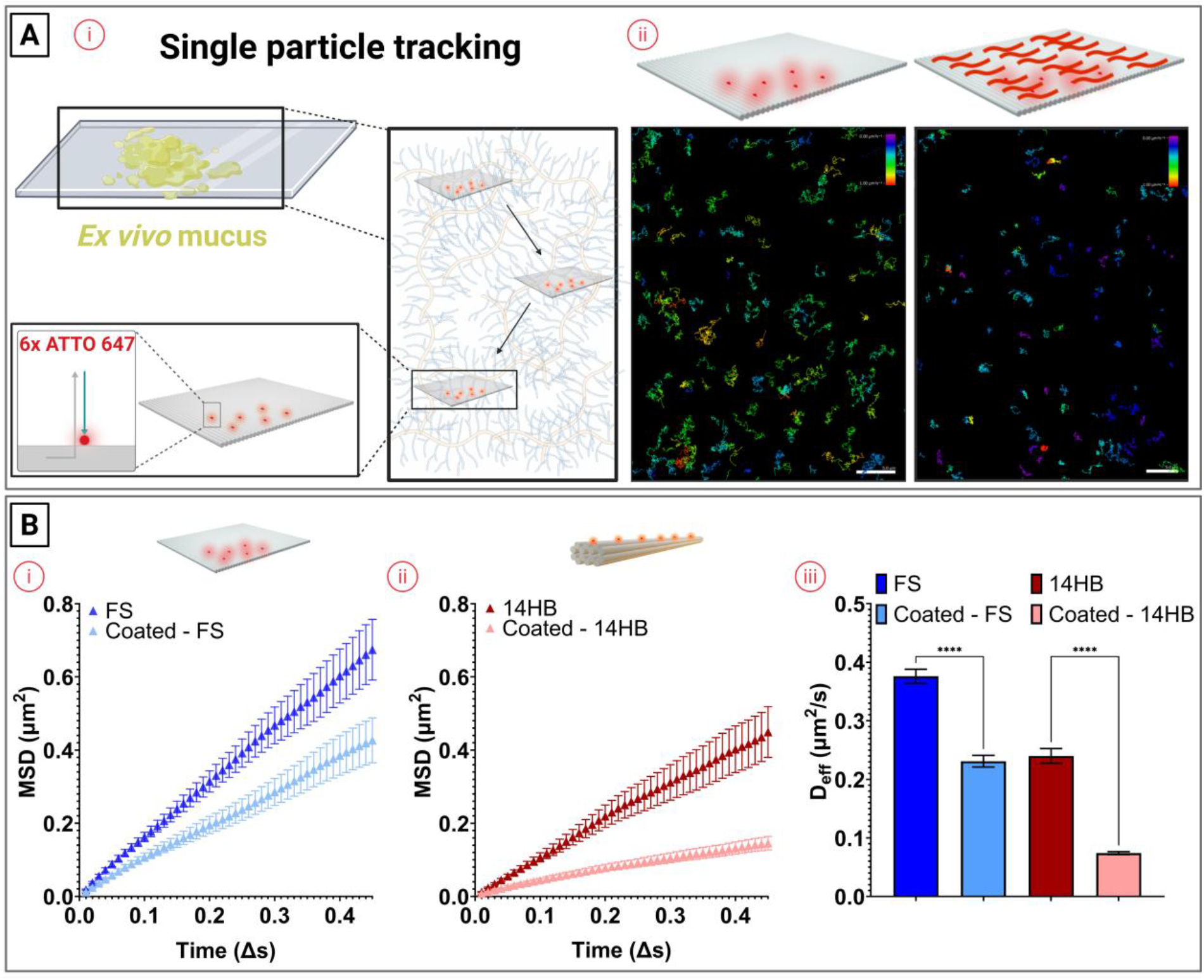
Single particle tracking of DNA origami in *ex vivo* intestinal mucus. (A) (i) Schematic illustrating the tracking of ATTO 647N-labeled DNA origami using HiLo microscopy; (ii) color-coded diffusion tracks of flat sheet origami with and without polymer coating (scale bar: 10 µm). (B) (i) Mean squared displacement (MSD) of flat sheet (FS) origami with and without coating; (ii) MSD of 14-helix bundle (14HB) origami with and without coating; (iii) diffusion coefficients of FS and 14HB, with and without coating. Data shown as mean ± SD (N = 100). Asterisks (****) indicate statistically significant differences (p ≤ 0.0001).

From these trajectories (N = 100 per condition), we computed the mean squared displacement (MSD), which describes the area covered by particle as a function of time and is a key metric to assess diffusive behavior.^45^ The MSD is calculated by averaging the squared distances that each particle travels over different time intervals. The shape of the MSD function indicates the type of motion dynamics. A higher slope of the MSD generally indicates greater particle mobility, corresponding to higher diffusion. Using the initial linear portion of the MSD vs. time lag plot, we extracted the effective Deff, which quantify how quickly a particle diffuses through the mucus matrix.^46^ This analysis allows us to directly compare the mobility of coated versus uncoated DNA origami structures and assess the impact of surface modification on mucus penetration.

For both DNA origami shapes (**Fig. 2B i and ii**), surface coating significantly reduced mucosal diffusion (p < 0.05), as evidenced by lower MSD and lower Deff. This effect is likely attributed to the positively charged nature of the coating polymer, which binds to both the DNA origami and mucin via electrostatic interactions. As previously reported, excessive positive surface charges, can hinder nanoparticle mobility by promoting adhesive interactions with the negatively charged mucin network, leading to particle immobilization within the mucus.^47–49^ Additionally, during tracking, we observed a tendency for coated particles to aggregate over time. This suggests that, while the polymer coating enhances the stability of the origami structures, it may also promote aggregation through electrostatic bridging with mucin, thereby impairing diffusion. When comparing the two shapes (**Fig. 2B iii**), the FS design consistently exhibited higher MSD and Deff (FS: 0.37 ± 0.02 µm^2^/s, 14HB: 0.21± 0.01 µm^2^/s), indicating superior mobility in mucus.^25^ Although polymer coating helped preserve the structural integrity and improve the stability of the DNA origami structures, it is not essential for mucosal penetration, as the origami remains stable in mucus for durations longer than typically required for effective transport.^19^ Based on these results, we selected the uncoated structures for subsequent experiments to investigate the impact of biomotor functionalization on mucus diffusivity. However, polymer coatings such as PCD may still be advantageous for downstream applications, particularly for enhancing cellular uptake and intracellular stability, making them relevant for future therapeutic strategies like siRNA delivery.^27,50,51^

### 2.3. Characterization of DNA origami functionalized with proteins

At low Reynolds numbers, where viscous forces dominate over inertial forces, creating geometrical asymmetry in synthetic motors is traditionally considered essential for achieving self-propulsion and enhanced diffusion.^52,53^ This is because an uneven or asymmetric shape enables directional movement in a highly viscous environment.^44,44^ However, recent studies have shown that an asymmetric molecular distribution, specifically, an uneven arrangement of surface-bound enzymes, can efficiently generate propulsion through localized catalytic activity.^32^ This observation motivated our use of DNA origami as a structural platform, owing to its programmability and ability to provide precise spatial control over enzyme positioning.

To investigate the influence of molecular asymmetry on nanomotor diffusivity, we designed two DNA origami nanostructures differing in the number and arrangement of single-stranded DNA (ssDNA) handles for site-specific enzyme attachment (**Fig. SI 5**). In the asymmetric configuration (Asym), five ssDNA extensions were positioned on one face of the origami. In the symmetric configuration (Sym), ten ssDNA handles were distributed across both edges of the nanostructure. This design strategy enabled a controlled comparison of enzyme spatial distribution on diffusivity.

For site-specific protein attachment, we first synthesized protein-DNA conjugates using either thiol-maleimide coupling or strain-promoted azide-alkyne cycloaddition (SPAAC) (**Fig. SI 6 and 7**). The resulting conjugates were then hybridized to the complementary ssDNA handles on the origami structure by simply mixing them at an excess relative to the number of binding sites.

To demonstrate the ability to achieve symmetric and asymmetric protein placement on the origami nanostructure, bovine serum albumin (BSA)-DNA conjugates were used (**Fig. SI 8**). Conjugation efficiency and structural integrity were assessed by agarose gel electrophoresis and AFM. A clear shift in the electrophoretic mobility of the DNA origami after conjugation indicated successful functionalization (**Fig. SI 8**). AFM imaging further verified BSA attachment, with protein-like ‘dots’ visible around the perimeter of the origami nanostructures. Although the Asym and Sym designs incorporated five and ten ssDNA extensions, respectively, full occupancy of all binding sites on individual structures was rarely observed, consistent with observations from previous studies.^27^ Nevertheless, the defined spatial arrangement of the available binding sites were sufficient to generate nanostructures with distinct protein distributions. This design strategy was guided by our initial BSA conjugation studies, where we observed that using an 8-fold excess of protein for the Sym structures and a 2×8-fold excess for the Asym structures yielded a similar average number of proteins per origami. However, the key distinction was in spatial localization: in the Asym configuration, proteins were concentrated on a single face of the structure, whereas in the Sym configuration, they were distributed bilaterally across both edges (**Fig. SI 8)**. These configurations thus provided a suitable platform for investigating how molecular asymmetry influences diffusive behavior.

### 2.4. Effect of urease on DNA origami diffusion

To enhance mucus penetration, urease-functionalized micromotors were employed to locally alter the microenvironment. Urease catalyzes the hydrolysis of urea into carbon dioxide and ammonia, resulting in a local pH increase (**Fig. 3**). This elevated pH induces a gel-to-sol transition in the mucus layer, thereby reducing its viscosity and facilitating enhanced nanoparticle diffusion.^8,32^

**Figure 3:**
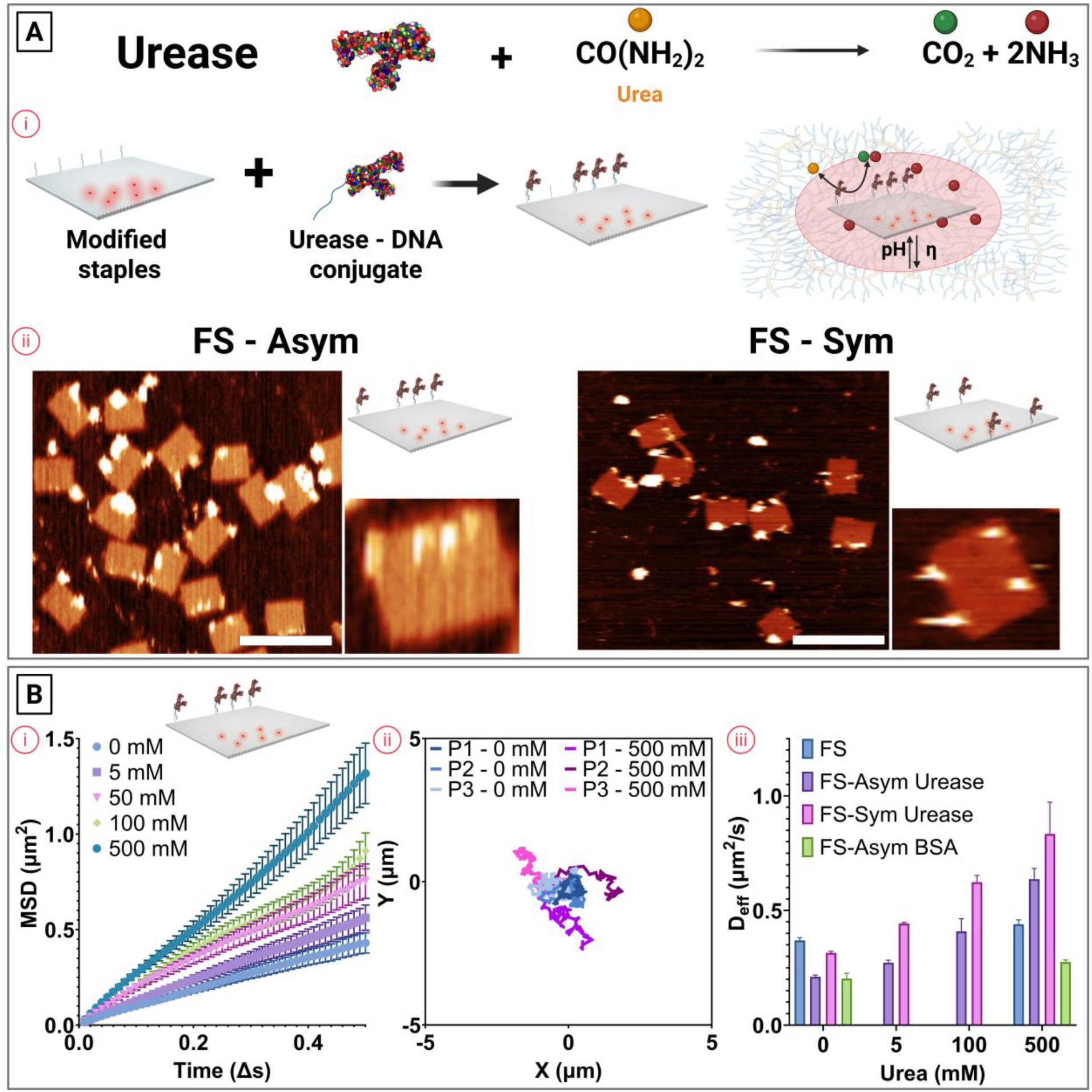
Single particle tracking of DNA origami-urease conjugates in *ex vivo* intestinal mucus. (A) Schematic representation of the enzymatic hydrolysis of urea by urease; (i) Schematic illustrating the conjugation of urease to DNA origami and the resulting local pH modulation, which alters the surrounding viscosity; (ii) Representative AFM images of Asym and Sym flat sheet (FS) urease-conjugated origami (scale bar: 200 nm). (B) (i) Mean squared displacement (MSD) of Asym-FS urease structures at varying urea concentrations; (ii) Representative diffusion tracks of Asym-FS urease structures at 0 and 500 mM urea; (iii) Diffusion coefficients of Asym- and Sym-FS urease conjugates compared to controls (naked FS and Asym-BSA) across different urea concentrations. Data are shown as mean ± SD (N = 100).

To assess the effect of urease on the diffusivity of DNA origami structures, urease-DNA conjugates were mixed with the FS DNA origami to generate the Asym and Sym configurations (**Fig. 3A i**). A clear shift in electrophoretic mobility was observed by agarose gel electrophoresis, indicating successful conjugation (**Fig. SI 9**). However, quantification of urease binding using AFM proved challenging, primarily due to the large molecular weight of the enzyme (~540 kDa), which limited the ability to accurately quantify the number of urease units per origami structure, as compared to BSA (**Fig. SI 9**). Despite this limitation, the combined evidence from gel shift assays and AFM imaging supports successful urease functionalization.^54^ Although precise stoichiometric determination remains difficult, the formation of Asym and Sym structures was consistently observed (**Fig. 3A ii**).

We first investigated the effect of urea concentration on the mobility of the DNA origami micromotors by measuring the MSD. As the concentration of urea increased, we observed a corresponding and significant rise in both MSD and Deff, indicating enhanced particle mobility (**Fig. 3B i**).^55^ This trend suggests that the catalytic activity of urease may contribute to particle motility, potentially by diffusiophoretic motion as well as local modification of the surrounding mucus. To further validate this, we extracted representative single-particle trajectories under fuel-free (0 mM urea) and high-fuel (500 mM urea) conditions. The trajectories clearly showed that particles exposed to urease and high urea concentrations exhibited greater displacement over time compared to those without fuel, confirming that enzymatic propulsion boosts diffusion through the mucus environment (**Fig. 3B ii**).

To confirm that effective propulsion requires direct conjugation of urease to the particles, we compared the diffusion behavior under identical conditions using (i) unmodified DNA origami in the presence of urease and urea in mucus, and (ii) BSA-coated particles with free urease and urea in mucus (**Fig. 3B iii**). In both cases, the enzymes were not bound to the DNA structures. These controls allowed us to evaluate whether enzyme activity alone, without surface conjugation, was sufficient to enhance particle diffusion. While a modest increase in diffusion was observed in both control conditions, the enhancement was significantly lower than that achieved with urease-conjugated particles. Specifically, at 500 mM urea, the Deff increased only slightly for the controls (1.21x for the naked FS and 1.36x for FS-Asym coated with BSA) compared to the baseline (0 mM). In contrast, particles with surface-conjugated urease exhibited much greater enhancement: a 3.02x increase for FS-Asym and a 2.65x increase for FS-Sym, clearly demonstrating the importance of enzyme conjugation for efficient propulsion.

Finally, when comparing the effect of enzyme distribution on the FS structure, either localized on one edge or evenly distributed on both edges, we observed that while the relative increase in diffusion was lower for the symmetric distribution, the absolute Deff was higher when enzymes were present on both edges. This suggests that for urease-powered DNA origami motors, a uniform distribution of enzymes enhances overall diffusion more effectively, potentially through a combination of diffusiophoretic motion and local modification of the microenvironment (e.g., changes in pH and viscosity).

### 2.5. Effect of catalase on DNA origami diffusion

To enhance mucus penetration, catalase-functionalized nanomotors were employed as an alternative strategy to urease-mediated diffusion enhancement. Urease and catalase can both influence particle motion through multiple mechanisms. Urease alters the local microenvironment by increasing pH and reducing mucus viscosity, while also promoting motion via diffusiophoresis. Catalase, on the other hand, can generate oxygen bubbles in confined environments and drive motion through diffusiophoretic effects (**Fig. 4**).^9^ In both cases, these mechanisms can act in combination, and their relative contribution depends on the concentration of the fuel. Specifically, catalase catalyzes the decomposition of hydrogen peroxide into water and oxygen, producing oxygen bubbles that propel the micromotors forward. This forms the basis of the hypothesis that bubble-driven propulsion offers an alternative strategy to overcome mucus barriers compared to urease-mediated mechanisms, which rely on local pH modulation to reduce mucus viscosity (**Fig. 4A i**).^9^ Catalase modification of the DNA origami was confirmed by gel electrophoresis (**Fig. SI 10**) and AFM (**Fig. 4A ii**), verifying successful conjugation. As with urease, it was challenging to accurately quantify the number of ligands per structure, but the formation of Asym and Sym structures was consistently observed.

**Figure 4:**
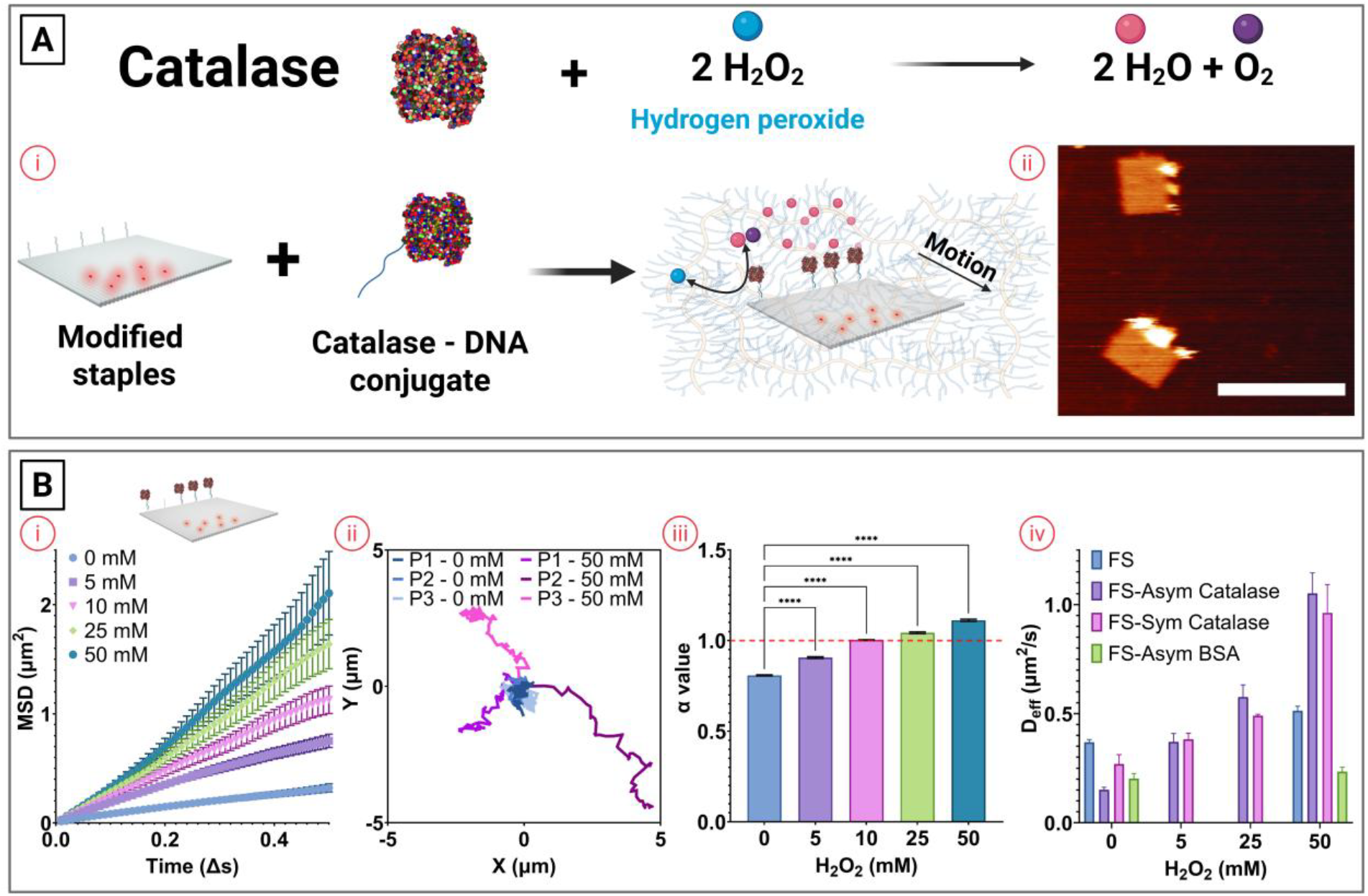
Single particle tracking of DNA origami-catalase conjugates in *ex vivo* intestinal mucus. (A) Schematic representation of the enzymatic breakdown of hydrogen peroxide by catalase; (i) Schematic illustrating the conjugation of catalase to DNA origami and the resulting generation of oxygen bubbles that can result in directed motion; (ii) Representative AFM images of Asym (FS) catalase-conjugated origami (scale bar: 200 nm). (B) (i) Mean squared displacement (MSD) of Asym-FS catalase structures at varying hydrogen peroxide concentrations; (ii) Representative diffusion tracks of Asym-FS catalase structures at 0 and 50 mM hydrogen peroxide; (iii) Diffusion coefficients of Asym- and Sym-FS catalase conjugates compared to controls (naked FS and Asym-BSA) across different hydrogen peroxide concentrations. Data are shown as mean ± SD (N = 100).

We then investigated the diffusion behavior of DNA origami structures asymmetrically functionalized with catalase across varying hydrogen peroxide concentrations (**Fig. 4B i**). As previously observed, the data indicated a clear concentration-dependent effect on diffusion.^9^ However, experiments were limited to a maximum concentration of 50 mM hydrogen peroxide, as higher concentrations (e.g. 100 mM) led to excessive bubble formation, causing substantial sample drift and rendering particle tracking unreliable (**Fig. SI 11**). When we extracted the particle tracks and calculated the anomalous diffusion exponent (α) for catalase-coated DNA origami, we observed signatures of directed motion (**Fig. 4B ii**). Although the motion was not consistently in a single direction, unlike externally guided systems such as light- or sound-driven nanomotors, we still observed signs of directed movement.^56,57^ Quantitative analysis of the α values further supported this observation (**Fig. 4B iii**). In diffusive systems, α ≈ 1 indicates Brownian motion, α < 1 suggests subdiffusion (often observed in mucosal diffusion due to confinement or crowding), and α > 1 reflects superdiffusion or active motion.^58^ At low hydrogen peroxide concentrations (e.g., 0 and 5 mM), the particles displayed subdiffusion (α < 1.0). However, at concentrations of 10 mM and above, α values consistently exceeded 1, reaching ~1.1, confirming a shift to enhanced, active transport. These findings are in line with the visual inspection of the tracks and support the hypothesis that catalase-driven propulsion can effectively promote self-mobility of nanoparticles through mucus.

Finally, when we compared the Deff of naked FS and FS-Asym particles coated with BSA in the presence of catalase and hydrogen peroxide in solution (i.e., without surface immobilization), we observed a similar trend to that seen with urease (**Fig. 4B iv**). Specifically, a significant enhancement in diffusion was only achieved when catalysis occurred in close proximity to the particle surface. In solution, the Deff increased modestly by 1.39x for naked FS and 1.15x for FS-Asym BSA, highlighting that free catalase in the surrounding medium has limited impact. These findings underscore the importance of surface-bound enzymatic activity in generating sufficient localized propulsion to enhance nanoparticle mobility.

Further analysis of the diffusion enhancement relative to the 0 mM hydrogen peroxide baseline showed a 6.95x increase in the Deff for FS-Asym catalase and a 3.57x increase for FS-Sym catalase. This marked difference indicates that, in contrast to the urease system, where symmetric and asymmetric enzyme distributions yielded comparable enhancements, catalase-driven propulsion is highly sensitive to surface symmetry. This can be attributed to the propulsion mechanism: catalase generates oxygen bubbles upon decomposition of hydrogen peroxide, and an asymmetric distribution of the enzyme results in a net directional thrust, leading to enhanced and more sustained motion. In the case of symmetric distribution, bubble generation occurs more uniformly around the particle, reducing net displacement and limiting the propulsion efficiency. These findings underscore the critical role of spatial ligand arrangement in bubble-propelled micromotors and suggest that optimizing particle asymmetry may be essential for maximizing diffusion in catalase-powered systems.

### 2.6. Time and shape effect on DNA origami diffusion

Finally, we sought to investigate how enzyme-driven propulsion influences mucosal diffusion over time (**Fig. 5A**), as well as how the origami shape and size impact transport efficiency within the mucus barrier (**Fig. 5B**).

**Figure 5:**
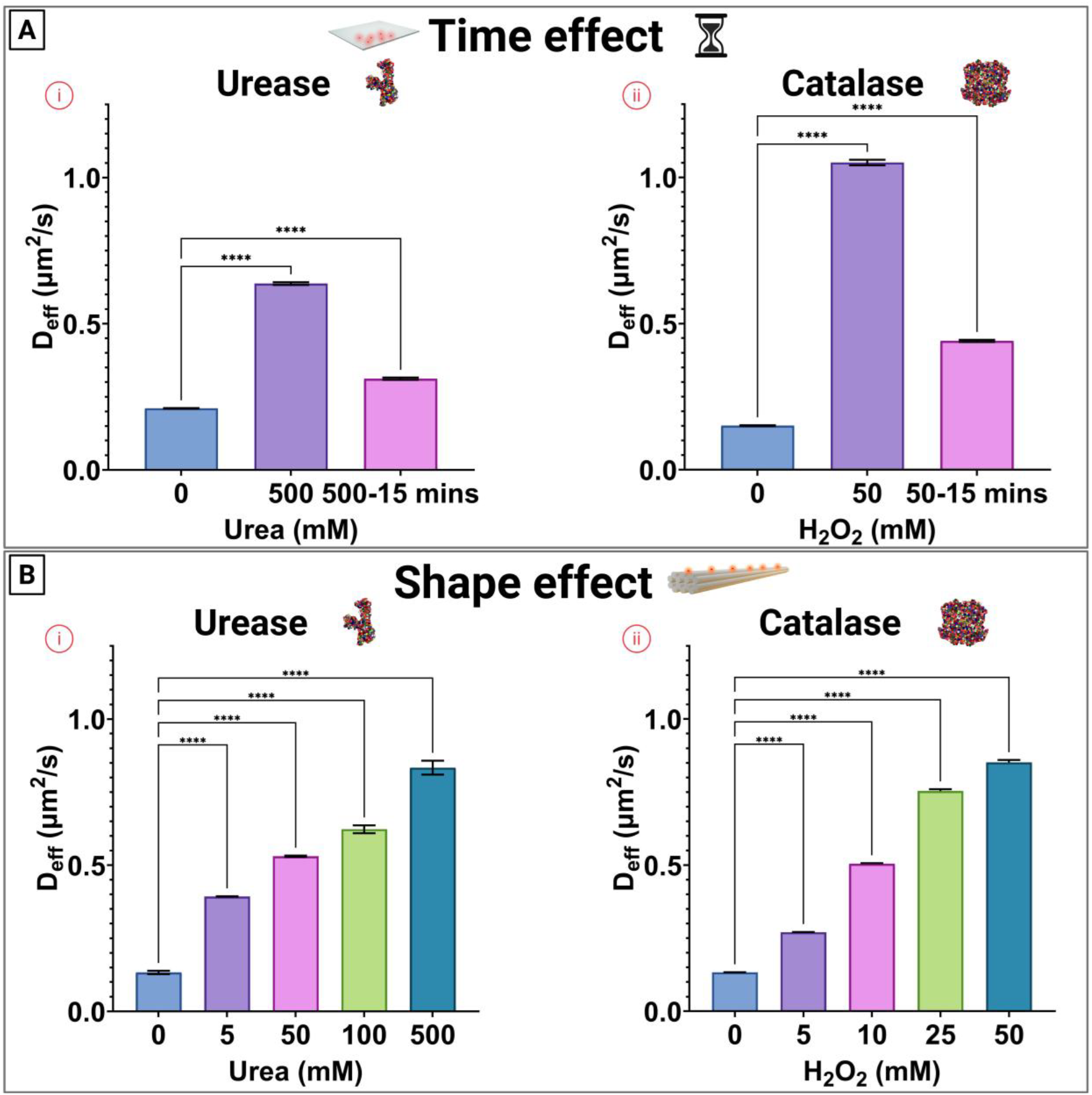
Time- and shape-dependent diffusion of DNA origami–enzyme conjugates in ex vivo intestinal mucus. (A): Diffusion coefficients of urease-(i) and catalase-conjugated (ii) origami over time in the presence of urea and hydrogen peroxide, respectively. (B) Diffusion coefficients of Asym-14HB origami conjugated to urease (i) and catalase (ii) at varying concentrations of urea and hydrogen peroxide. Data are presented as mean ± SD (N = 100).

We measured the Deff of the particles 15 minutes after initial incubation (**Fig. 5A**). Although the Deff had decreased by approximately 50% compared to its peak value, we still observed enhanced motion for both urease-(**Fig. 5A i**) and catalase-coated (**Fig. 5A ii**) particles. This suggests that enzyme-driven propulsion can sustain enhanced particle mobility over physiologically relevant timescales. Considering that mucus turnover in the small intestine occurs over tens of minutes to a few hours, this prolonged activity window indicates that fuel-driven nanocarriers could maintain sufficient mobility to penetrate the mucus, and potentially reach the epithelial barrier before being cleared.^19^

To check the effect of different shapes, we employed a 14HB DNA origami structure, an elongated rod-like geometry approximately 155 nm in length and asymmetrically conjugated either urease or catalase to its surface (**Fig. 5B**). We then evaluated their Deff across a range of urea (**Fig. 5B i**) and hydrogen peroxide (**Fig. 5B ii**) concentrations. Consistent with previous findings using FS structures, increasing fuel concentration resulted in higher Deff values for both propulsion systems, confirming the activity-dependent enhancement of mobility. However, a key difference emerged when comparing the propulsion efficiencies between the two shapes. While catalase-coated FS structures had previously outperformed urease-coated ones at similar conditions, the 14HB structures displayed comparable Deff values for both enzyme systems at their respective optimal fuel concentrations. This observation suggests that, despite successful enzyme-driven enhancement of mobility, the intrinsic size of the 14HB structures (~155 nm) may approach a critical threshold in relation to the mucus mesh pore size (typically 150–200 nm).^20,37^ In such cases, steric hindrance becomes a limiting factor that may restrict further diffusion gains, even in the presence of active propulsion. This contrasts with smaller FS structures, which can more readily take advantage of propulsion mechanisms to navigate through the mucus network.^20^

These findings are in line with our previous work, where we demonstrated that asymmetric ligand distribution, specifically charge anisotropy, could significantly enhance nanoparticle diffusion through mucus.^25^ While direct 1:1 comparisons between structures of different shapes are complicated by differences in scaffold and experimental conditions (e.g., *ex vivo* mucus heterogeneity), our current results further support the hypothesis that shape constraints, in addition to surface properties and propulsion mechanisms, play a critical role in dictating the diffusion performance of nanoscale drug carriers in mucosal environments.^37^ Although enzyme-mediated propulsion (whether via urease or catalase) clearly enhances mobility of the structures, we observed that such active motion may still be fundamentally limited by the physical barrier imposed by the mucus mesh. In particular, when the particle dimension approaches or exceeds the average mucus pore size (~150-200 nm), even active transport mechanisms may no longer overcome the steric hindrance. This highlights the need for nanoparticle designs that account for surface properties, propulsion efficiency, and biophysical compatibility with the target barrier to optimize delivery performance.

## 3. Conclusion

We have explored the properties and potential of DNA origami in *ex vivo* mucus and biological fluids, where we have employed FRET-based analysis to confirm and quantify the structural stability of the nanostructures in biologically relevant environments such as porcine intestinal fluid and mucus over extended time periods. These studies demonstrate that DNA origami maintains its integrity in complex, biologically relevant environments such as porcine intestinal fluid and mucus, highlighting its promise for applications in oral drug delivery and as a platform for investigating particle–mucus interactions under physiologically relevant conditions.

To gain mechanistic insights into enzyme-driven motion, we used SPT to assess how different bio-enzymes and their spatial distribution influence diffusion behavior. Our results showed that for both urease and catalase, enhanced diffusion was only observed when the enzymes were anchored to the DNA origami surface; the presence of enzymes in the mucus medium was insufficient to induce significant enhanced motion. Importantly, we demonstrated that this enhanced propulsion was maintained for tens of minutes, rather than occurring only for a few seconds.

Building on our initial mechanistic studies of urease spatial arrangement and anisotropy distribution on DNA origami, this work extends the investigation to a biological setting in mucus, examining both spatial distribution and enzyme type. We also investigated the impact of spatial asymmetry in enzyme placement and found enzyme-specific effects on diffusion. For urease, symmetrically distributed enzymes led to improved diffusion, likely due to a more uniform local increase in pH and reduction in mucus viscosity. In contrast, for catalase, an asymmetric distribution with enzymes localized on one edge of the origami structure resulted in the highest diffusion rates, likely because bubble generation on a single edge enables more directional propulsion, whereas symmetric placement produces opposing thrusts that reduce overall efficiency.

Finally, we assessed the role of particle shape, particularly in relation to the mucus mesh size. Increasing origami length did lead to enhanced motion for both enzyme systems, but the effect was most likely limited by steric hindrance within the mucus network.

Overall, this work reaffirms the importance of nanostructure shape and ligand distribution for navigating mucosal barriers and introduces enzyme propulsion modality and spatial organization as additional, tunable parameters for optimizing diffusion. These findings highlight the potential of pairing enzyme-powered DNA origami systems with therapeutics, particularly in pathophysiological conditions where endogenous fuels are elevated, opening new avenues for targeted mucosal drug delivery.

## 4. Materials and Methods

### Materials

All chemicals were used as received unless otherwise stated. The ingle-stranded M13p18 scaffold for both the flat sheet (FS) and 14-helix bundle was purchased from Tilibit, while staple strands, handle-extended staple strands, modified DNA oligonucleotides, and fluorescently modified DNA oligonucleotides, were obtained from Integrated DNA Technologies. DNase I was purchased from Thermo Fisher Scientific. All other chemicals and reagents, including: DBCO-PEG4-NHS ester, BSA, catalase, urease, phosphate-buffered saline (PBS), glycogen, dimethyl sulfoxide (DMSO), NaCl, sodium tetraborate, DMF, MgCl2, CaCl2, HEPES, ethanol, NaOAc, EDTA, RNase-free MQ were purchased from Sigma-Aldrich (St. Louis, MO). Solvents were HPLC grade and used as received. Ultrapure water was used throughout the experiments and obtained from Merck Millipore Q-Pod system (Merck Group, Burlington (MA), U.S.A) with a 0.22 μm Millipore Express 40 filter (18.2 MΩ).

### Data and statistical analysis

Data analysis was performed with GraphPad Prism (Version 9.4.1 (681) Insight Partners, Graphpad Holdings, LLC, New York City (NY), U.S.A.). Data are presented as average and standard derivation (SD) or standard error measurements (SEM) where n represents the number of repetitions within each sample and N represents the number of samples. Statistical analysis was performed for statistically significant differences (*p < 0.05) with GraphPad Prism using a t-test (for two independent populations) or one-way analysis of variance (ANOVA) (three or more independent populations). The figures were created with Biorender.com. Image analysis and processing was done using the software ImageJ (version 1.53t, National Institutes of Health, USA).

### Methods

#### Design and assembly of DNA origami

The design of the FS origami and 14-helix bundle was designed using the cadnano software;^**59**^ further details are outlined in the **Fig. S1 and S3**, respectively. The structures were assembled by using a single-stranded M13p18 scaffold at concentrations ranging from 3-40 nM, accompanied by a 10-fold molar excess of the staple strands. The sequences of all oligonucleotides used are provided in **Tables SI 1-4** for the FS origami, and **Tables SI 5-7** for the 14-helix bundle. The folding buffer contained 20 mM MgCl2, 50 mM NaCl, and 1x TAE. The FS origami was annealed by using the following method: 95 ^°^C for 5 minutes, 80 ^°^C for 2 minutes, 80 ^°^C for 3 minutes, 80-65 ^°^C ramp (0.5 ^°^C/2 min), 55-25 ^°^C ramp (0.3 ^°^C/48 secs) and 4 ^°^C hold. The 14-helix bundles were annealed by using the following method: 95 ^°^C for 1 minute, 70 ^°^C for 3 minutes, 70-40 ^°^C ramp (1 ^°^C/30 min) and 20 ^°^C hold.

#### Synthesis of fluorophore-modified DNA conjugates

Amino-modified DNA was mixed with 0.1 mg Cy3- or Cy5-NHS ester (Sigma-Aldrich), sodium tetraborate (0.1 M, 15 µL pH 8.5) and 15 µL DMF. The sample was then incubated overnight on a shaker at room temperature. After, the solution was diluted to 100 µL with MQ and then mixed with 250 µL 96% EtOH, 14 µL NaOAc (3 M, pH 5.2) and 1 µL glycogen (20 mg/ml). The mixture was flash frozen and centrifuged (14,000 rpm) at 4 °C for 45 min. Lastly, the pellet was resuspended in MQ and purified by RP-HPLC with Phenomenex Clarity 3u Oligo-RP 50 mm × 4.6 mm column running a gradient of acetonitrile in TEAA buffer (0.1 M, pH 7). The fractions containing product were collected and lyophilized before being dissolved in MQ. The strand modified with ATTO 647N was directly purchased.

#### Mucus isolation

Intestines from healthy fasted (18–24 h) gilts (40–60 kg, 3–4 months, Danish Landrace) were obtained after experimental surgery. Immediately after euthanization, up to 5 m jejunum was isolated distal to the ligament of Treitz. Sections were opened by a latitude cut and porcine intestinal mucus was isolated by gently scraping the mucosal surface. Mucus was kept on ice at all times and stored at −20 °C until use. Procedures were according to the authorization by Danish Veterinary and Food Administration (license number: 2020-15-0201-00610).

#### FRET experiments

The FS origami was annealed and purified from the excess staples using 100 kDa Amicon Ultra-0.5 mL centrifugal filters (Millipore, Germany) into a high salt buffer (12.5 mM MgCl2, 50 mM NaCl, and 1x TAE). The structures were then mixed with a 5-fold molar excess per binding site of the Cy3- and Cy5-modified DNA strands and left to incubate overnight at room temperature. Next, the structures were purified from the excess fluorophore strands using 100 kDa Amicon Ultra-0.5 mL centrifugal filters, either into a high salt buffer for the controls (12.5 mM MgCl2, 50 mM NaCl, and 1x TAE), or a low salt buffer (0.6 mM MgCl2, 0.5 mM CaCl2, 10 mM HEPES, pH 7.4). The samples washed in low-salt buffer were subsequently mixed with the protective cationic polymer poly(cystaminebisacrylamide-1,6-diaminohexane) (PCD), synthesized in-house, (0.56 μL of 5 mg/mL per 10 nM, 10 μL of the origami) and left to incubate for 1 hour before use.^27^ DNA-origami samples were prepared at 50 nM in folding buffer and diluted 1:10 immediately before measurement to give a final concentration of 5 nM in the test medium. For each condition, 10 µL of the 50 nM origami solution were mixed with 90 µL of either (i) ex vivo porcine intestinal mucus, (ii) ex vivo porcine intestinal fluid, or (iii) high-salt buffer (positive control) and iv) DNase 10 units/mL (negative control) in a flat-bottom 96-well plate. To minimize evaporation during the 14 h acquisition, each well was sealed with an optically clear film. Fluorescence spectra were recorded on a plate reader (Tecan Spark, TecanGroup Ltd., Männedorf, CH) equilibrated to 37 °C (excitation = 530 nm; 555–705 nm emission scan, 1 nm steps). Donor and acceptor intensities were taken at their respective emission maxima (Cy3 ≈ 565 nm, Cy5 ≈ 665 nm). To account for slight well-to-well variations and the high-salt control, each spectrum was area-normalized to unity before analysis. Ratiometric FRET efficiency (E) was then calculated for every time point using the sensitized-emission equation:

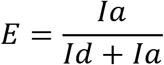

where Ia is the acceptor emission upon donor excitation and Id is the donor emission under the same conditions. Normalized E values were plotted as a function of time to compare structural stability (N=3).

#### BSA-DNA conjugation reaction

To produce the maleimide-modified DNA strand, amino-modified DNA (10 nmol in 50 μL of MQ) was mixed with 4-(N-maleimidomethyl) cyclohexanecarboxylic acid N-hydroxysuccinimide ester (SMCC, 598 nmol in 50 μL of dry DMF) and 0.25 μL of TEA. The sample was then incubated overnight on a shaker at room temperature. After, the solution was then mixed with 250 μL 96% EtOH, 14 μL NaOAc (3 M, pH 5.2) and 1 μL glycogen (20 mg/ml). The mixture was flash frozen and centrifuged (14,000 rpm) at 4 ^°^C for 45 min. Lastly, the pellet was resuspended in MQ and purified by RP-HPLC with Phenomenex Clarity 3u Oligo-RP 50 mm × 4.6 mm column running a gradient of acetonitrile in TEAA buffer (0.1 M, pH 7). The fractions containing product were collected and lyophilized before being dissolved in MQ and stored at −20^°^C until required. For optimizing the conjugation between the maleimide-DNA strand and BSA, the protein (0.5 nmol) was mixed in buffer (10 mL of 100 mM sodium phosphate supplemented with 550 mM NaCl, with varying pH), with varying equivalence of the maleimide-DNA strand. The reaction was left overnight at room temperature or at 37 °C. The samples were mixed with 2.5 mL NuPAGE^TM^ LDS Sample Buffer (Invitrogen) and heated to 70 °C for 10 min before running on a NuPAGE^TM^ 4-12% Bis-Tris polyacrylamide gel (Invitrogen) at 200V for 1 hour. For the ladder, 4 mL SeeBlue^TM^ Plus2 Prestained Standard (Invitrogen) was used. The gels were then stained with SimplyBlue SafeStain (Life Technologies) and visualization with a Gel Doc EZ (Bio-rad). From these optimizations, using a 10-fold molar excess of the DNA strand, with a buffer at pH 7.2 for an overnight reaction resulted in the highest conversion yield. These conditions were used for an upscaled reaction of 200 mL. After, the mono-labeled BSA-DNA conjugate was purified using HPLC with a DNAPac^TM^ PA-100 BioLC 4 × 250 nm column (Thermo Scientific) running a gradient of 25 mM TRIS supplemented with 1M NaCl, pH 8, followed by a gradient of 25 mM TRIS, pH 8. After isolation, the conjugates were concentrated using 30 kDa Amicon Ultra-0.5 mL centrifugal filters into a buffer of 1x PBS. The concentration was calculated using the extinction coefficient of 43,824 M^−1^cm^−1^ at 279 nm.

#### Enzyme-DNA conjugation reaction

For the enzyme-DNA conjugates, the urease and catalase (5 nmol in 1x PBS) were independently mixed with DBCO-PEG4-NHS ester (40 nmol in dry DMSO) to reach a final reaction volume of 1 mL. The buffer used for the reaction was 1x PBS, with 60 μL of DMSO). The samples were left to incubate overnight at 4 °C whilst shaking at 350 RPM. The constructs were then purified from the excess DBCO using 100 kDa Amicon Ultra-0.5 mL centrifugal filters in 1x PBS. The concentration of the product was calculated using the extinction coefficient of 54,165 M^-1^cm^-1^ at 280 nm for urease and 246,000 cm^-1^M^-1^ at 280 nm for catalase. The enzyme-DBCO conjugates were then mixed with varying molar equivalence of the DNA-azide strand in 1x PBS and left to incubate overnight at 37 °C. The conjugates were analyzed using SDS-PAGE, as described in the BSA-DNA conjugate section. From these optimizations, using a 1-fold molar excess of the DNA strand resulted in the highest conversion yield for the urease-DNA conjugates, while a 5-fold molar excess was optimal for the catalase-DNA conjugates. These conditions were used for an upscaled reaction of 600 μL. The conjugates were purified using 30 kDa Amicon Ultra-0.5 mL centrifugal filters.

#### Assembly and visualization of the origami-protein conjugates

The purified origami structures, either with zero, five, or ten protein capture extensions, were mixed with varying molar equivalences per binding site and left to incubate overnight at room temperature. A 1% agarose gel containing 12.5 mM MgCl2 and 1x TBE was used to analyze the binding of the protein-DNA conjugates to the origami structures. The gels were eluted at 65 V for 90 min and pre-stained with 1x SYBR Safe (Invitrogen). The bands of interest were excised, and the solution was extracted by squeezing the gel between two glass slides. The samples were used for AFM or TEM.

#### AFM

Freshly cleaved mica was incubated with NiCl2 (200 mM, 50 μL) for 2 min before being blotted and washed with 50 μL MQ. The droplet was dried with nitrogen gas and the unpurified sample (1 nM, 5 μL) was immediately deposited onto the mica surface and incubated for 2 min. 50 μL of folding buffer was deposited both on the mica and the liquid cell. The sample was scanned using an Olympus BioLever-mini cantilever (spring constant 0.09 N m^-1^, 25 kHz resonance frequency in water) in liquid tapping mode on a Bruker Multimode-8 AFM. All images were analyzed using Gwyddion analysis software.

#### TEM

The purified sample (5 μL, 6 nM) was deposited onto a glow-discharged carbon-coated grid (400 mesh, Ted Pella) for 3 min. The grid was then blotted, dipped on 8 μL of MQ before being blotted again. The grid was immediately treated twice with 2% uranyl formate (4 μL) with a 20 second hold before blotting the second treatment. The grids were left to dry for 2 min. Imaging was conducted with a Tecnai G2 Spirit TEM, operated at 120 kV.

#### Single Particle Tracking

The origami structures were annealed, either with a 10-fold excess of zero, five (Asym), or ten (Sym) protein capture extensions, and a 10-fold excess of the six ATTO 647N-DNA capture extensions. Additionally, a 25-fold molar excess of the ATTO 647N-modified DNA strand was used. After annealing, the structures were purified from the excess staple strands and excess fluorophore strands using 100 kDa Amicon Ultra-0.5 mL centrifugal filters. The high salt buffer was used during the washing steps. The structures were then diluted to 10 nM and mixed with an 8-fold molar excess of the protein-DNA conjugates and left to incubate overnight at room temperature. DNA-origami samples were prepared at 10 nM in folding buffer and diluted 1:10 immediately before measurement to give a final concentration of 1 nM. Images were acquired in a Nanoimager® (ONI, Oxford) using the NimOS software with a 640 nm laser (190 mW). The sample was illuminated using a highly inclined and laminated optical sheet (HiLo) at an angle of 47°, at 15% laser power. Fluorescence was recorded using an ONI 100x, 1.49 NA oil immersion objective and passed through a quad-band pass dichroic filter. Images were acquired onto a 425×518-pixel region (pixel size 0.117 μm) for 10 s at 100 fps. For each experimental condition, a total of at least 3 different biological samples were analyzed. The results were filtered using the NimOS software with the following settings: maximum frame gap = 5, minimum distance between frames = 0.800 µm, exclusion radius = 1.200 µm, a minimum number of steps = 50, a minimum diffusion coefficient of 0.01 μm^2^s^−1^. With these parameters, track steps were extracted, and analyzed using a Python-based code1 to obtain the trajectories of the DNA origami (N = 100) and calculate the mean-squared displacement (MSD) with the following equation (2):

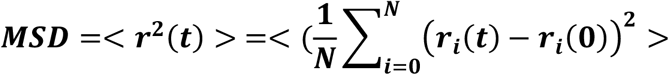

Then, the diffusion coefficient was obtained by fitting the MSD data to equation (3), where r = radius and t = sampling time and MSD(t) = 2dD, where D = diffusion coefficient and d = dimensionality (ONI measurements have dimension d = 2):

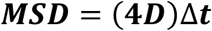

was used to fit the MSD curves.

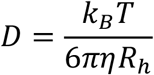

where *D* = diffusion coefficient, kB = Boltzmann constant, *T* = temperature, *η* = mucus viscosity and Rh = hydrodynamic radius.^60^

To extract the α value, the MSD was plotted against lag time on a log–log scale, and a linear fit was applied; the slope of this fit corresponds to the anomalous diffusion exponent α.

## CRediT authorship contribution statement

**Matteo Tollemeto** (Conceptualization; Data curation; Formal analysis; Investigation; Methodology; Validation; Visualization; Writing – original draft; Writing – review & editing) **Emily Tsang** (Data curation; Formal analysis; Investigation; Methodology; Validation; Visualization; Writing – review & editing) **Lars J.M.M. Paffen** (Methodology; Writing – review & editing) **Lasse Højlund Eklund Thamdrup** (Supervision; Writing – review & editing) **Jan van Hest** (Conceptualization; Supervision; Writing – review & editing;) **Tania Patiño Padial** (Conceptualization; Supervision; Writing – review & editing;) **Kurt V. Gothelf** (Conceptualization; Supervision; Writing – review & editing; Project administration; Funding acquisition) **Anja Boisen** (Conceptualization; Supervision; Writing – review & editing; Project administration; Funding acquisition)

## Acknowledgements

The authors would like to acknowledge the Danish National Research Foundation (DNRF122) and Villum Fonden (Grant No. 9301) for intelligent drug delivery and sensing using microcontainers and nanomechanics (IDUN), the Novo Nordisk Foundation (CEMBID: NNF17OC0028070) and (NNF17OC0026910) and the European Research Council in Foldable, REconfigurable & Jagged devices for enhanced drug Absorption/seeding (FREJA), grant ref. No. 101054945.

The authors would also like to acknowledge Prof. Jørgen Kjems and the Bioimaging Core Facility, Health, Aarhus University, Denmark, for providing access to equipment and support with the ONI system.

