## Supplementary information for "Enzyme-powered DNA origami nanostructures for enhanced mucosal diffusion"

**Corresponding authors:**

**Supplementary Figure 1. Blueprint of the scaffold and staple routing for the flat sheet origami.** Illustration of the scaffold and staple routing for the flat sheet origami. This visual representation showcases the routing of both scaffold and staple strands, exported from Cadnano.

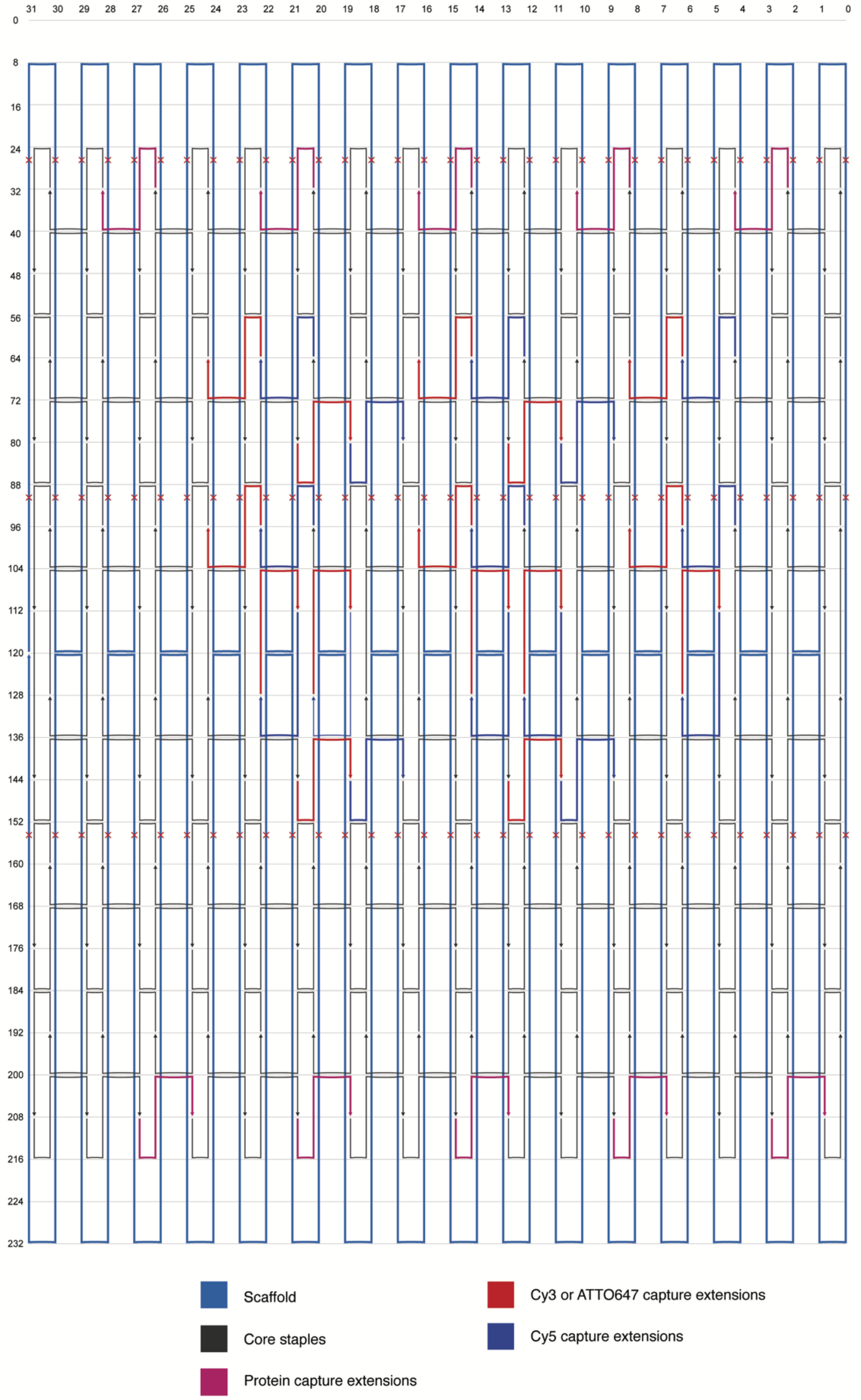

**Supplementary Figure 2. AFM image of the unfunctionalized flat sheet origami.**

Full AFM image of the unfunctionalized flat sheet origami. Before imaging, the structures were purified from the excess staples via Amicon filtration in a buffer containing 12.5 mM $MgCl_2$, 50 mM NaCl, 1x TAE.

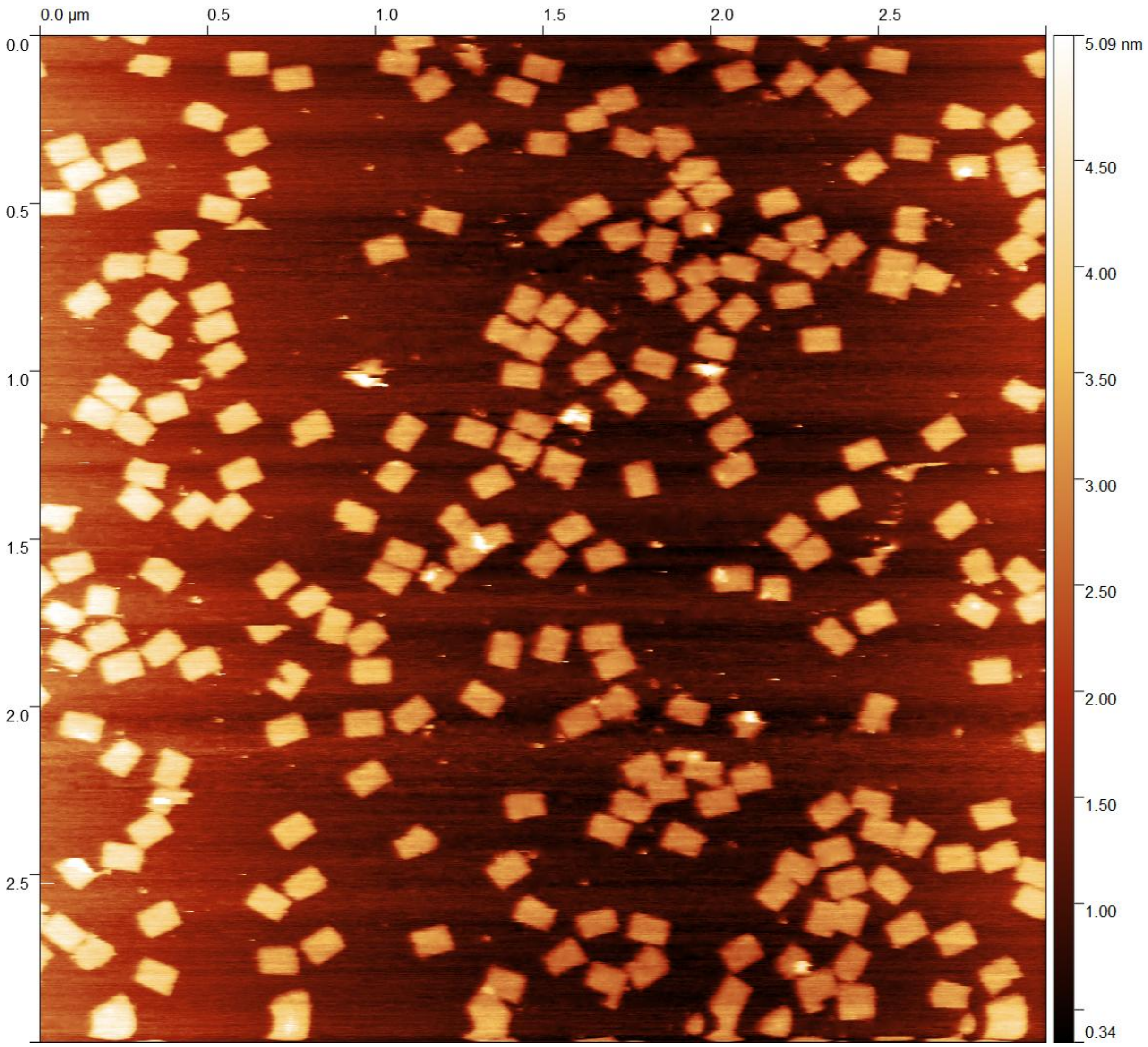

**Supplementary Figure 3. Design schematic and strand for the 14-helix bundle (14HB) origami.**
(a) Illustration of the scaffold and staple routing for the 14HB origami, as exported from Cadnano. (b) Color scheme indicating the different strands used in the assembly of the 14HB structure. (c) Front view of a simplified schematic showing the positions of strand extensions. Two designs variants were used. (d) A design incorporating five protein capture strands positioned on one edge of the 14HB, enabling a theoretical maximum of five protein attachments per origami (Asym configuration); (e) A design with ten protein capture strands distributed on both edge of the 14HB, allowing for a theoretical maximum of ten protein attachments per origami (Sym configuration).

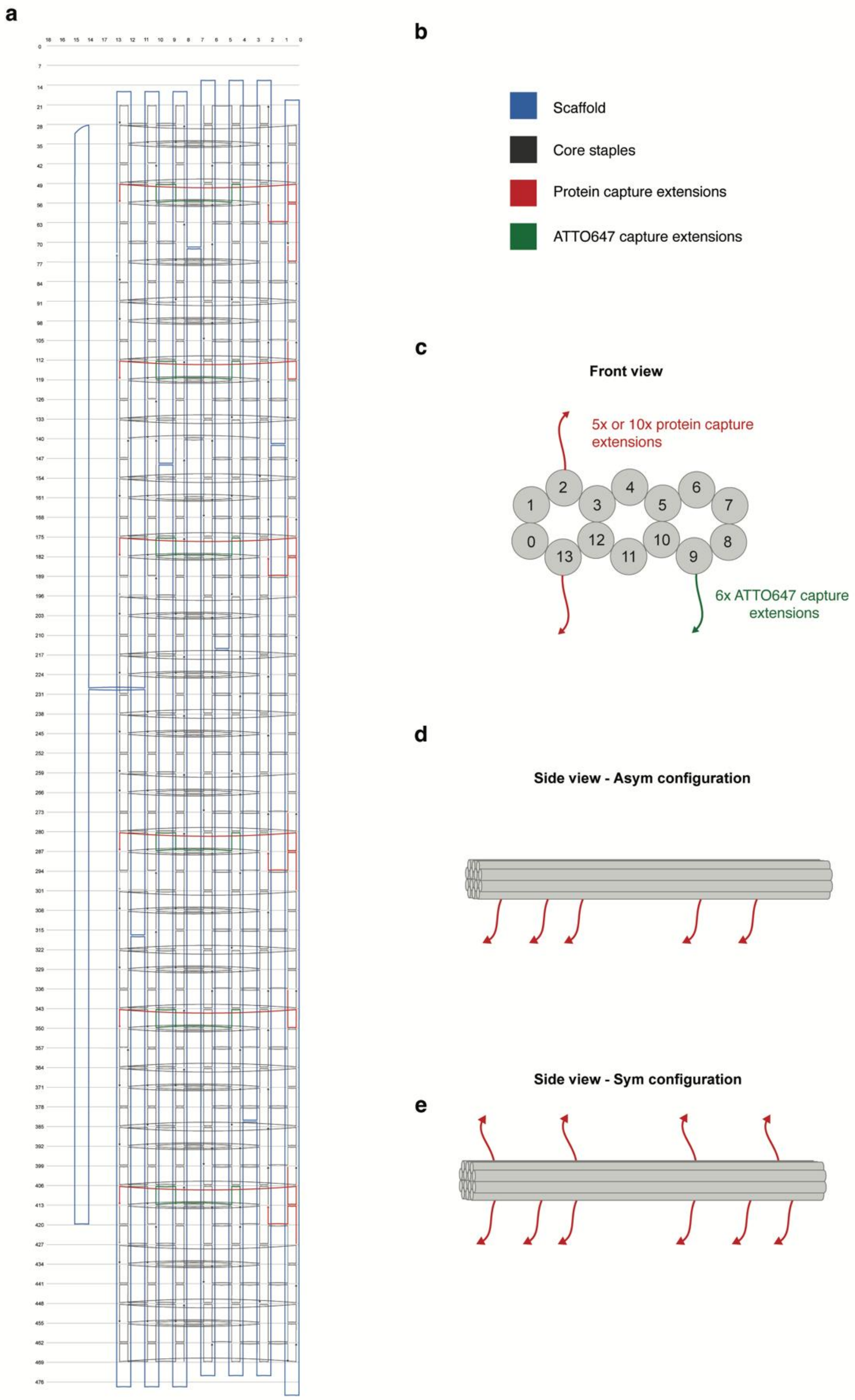

**Supplementary Figure 4. TEM image of the unfunctionalized 14HB origami.**

Full Cryo-TEM image of the unfunctionalized 14HB origami. Before imaging, the structures were purified from the excess staples via Amicon filtration in a buffer containing 12.5 mM $MgCl_2$, 50 mM NaCl, 1x TAE. Scale bar: 200 nm, magnification: 30,000x.

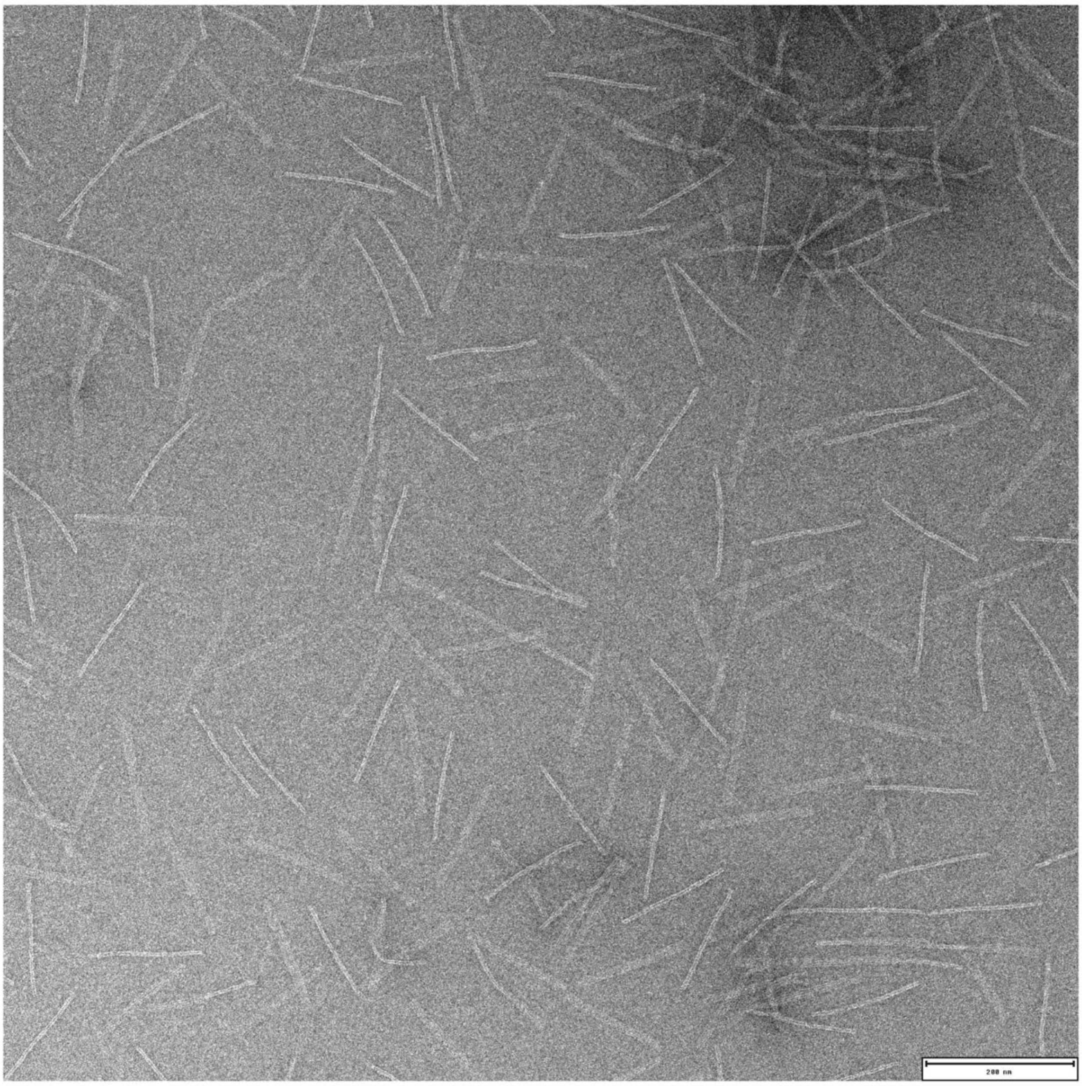

**Supplementary Figure 5. Schematic overview of protein-flat sheet DNA origami conjugates.**
Simplified schematic illustrating the conjugation strategy used to attach protein-DNA conjugates to flat sheet DNA origami structures. Two distinct designs were used: one with five protein capture strands located on one edge of the flat sheet, allowing for a theoretical maximum of five proteins per origami (Asym configuration); and another with ten protein capture strands distributed on both edges of the flat sheet, allowing for a theoretical maximum of ten proteins per origami (Sym configuration).

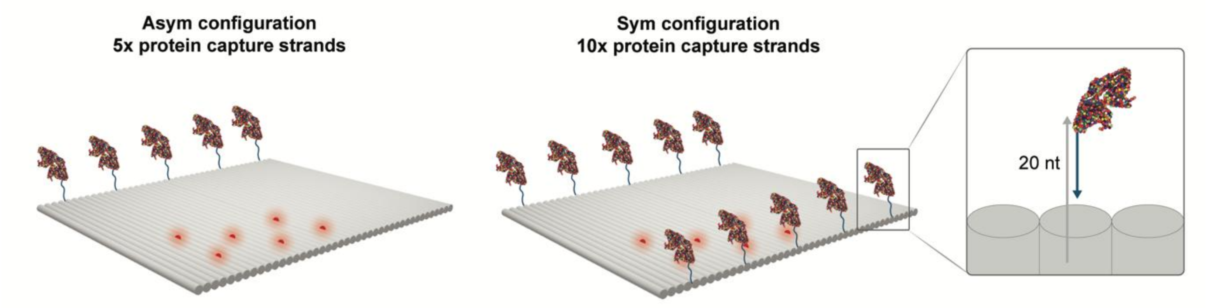

**Supplementary Figure 6. Bioconjugation of bovine serum albumin with DNA.**
(a) Reaction scheme for the conjugation of an amino-modified DNA strand with sulfosuccinimidyl 4-(N-maleimidomethyl)cyclohexane-1-carboxylate (sulfo-SMCC) via a NHS ester functional group. (b-c) Optimization of the bioconjugation conditions for the reaction between maleimide-functionalized DNA and bovine serum albumin (BSA), examining the effects of buffer pH, DNA-to-protein molar ratio, and reaction temperature. All reactions were incubated overnight then ran on a 4-12% polyacrylamide gel and post-stained with Coomassie blue to visualize the proteins present in the gel. The BSA-DNA conjugate is indicated with a black arrow, and the corresponding band intensity, which is used as a proxy for conjugation efficiency, is annotated above the product band.

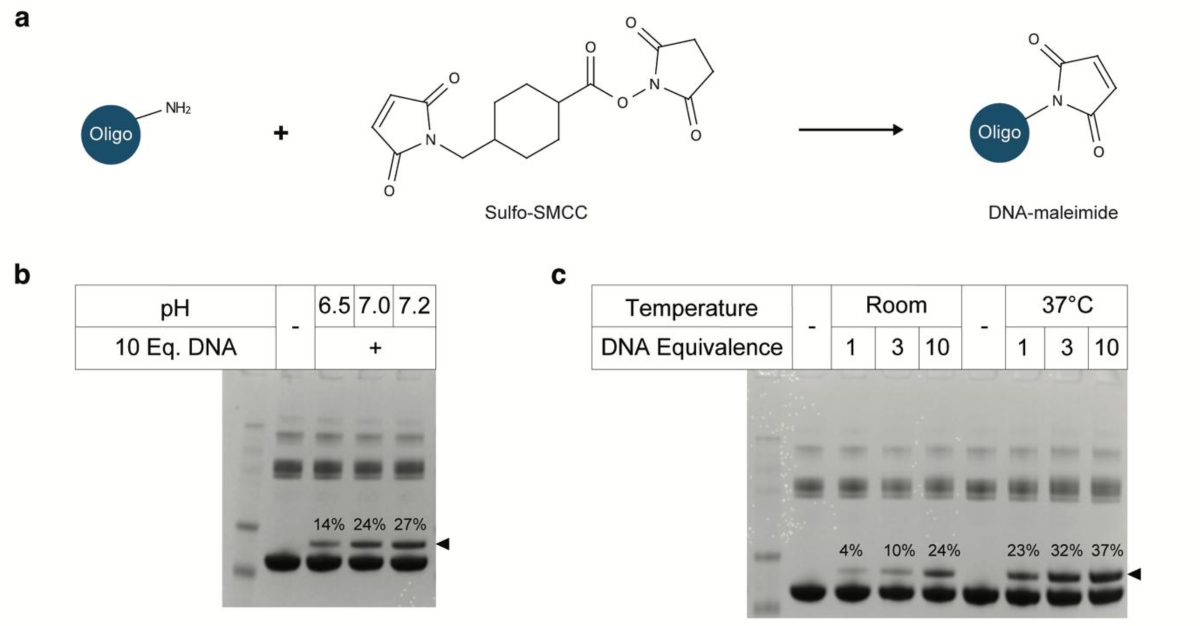

As BSA typically contains only one free cysteine residue in the reduced state, the resulting bioconjugate was pre-dominantly mono-labeled with a single DNA strand. The final optimized conditions used in subsequent reactions consisted of a buffer at pH 7.2, a 10-fold molar excess of maleimide-DNA, and an incubation time of overnight at 37°C.

**Supplementary Figure 7. Bioconjugation of urease and catalase with DNA.**
Schematic representation of the two-step conjugation of DNA to urease and catalase. (a) The enzyme was first reacted with dibenzocyclooctyne-N-hydroxysuccinimidyl ester (DBCO-NHS ester), followed by purification using Amicon filtration. (b) The resulting enzyme-DBCO conjugate was then incubated with an azide-modified DNA strand to yield the enzyme-DNA conjugate. Optimization of the DNA-to-protein molar ratio was carried out for both enzymes. All reactions were incubated overnight at 37°C and analyzed by 4-12% polyacrylamide gel electrophoresis, followed by Coomassie blue staining to visualize the proteins. (c, d) Gel electrophoresis results for urease and catalase, respectively. Black arrows indicate the enzyme-DNA conjugates corresponding to different values of *n*, representing the number of DNA strands conjugated per protein.

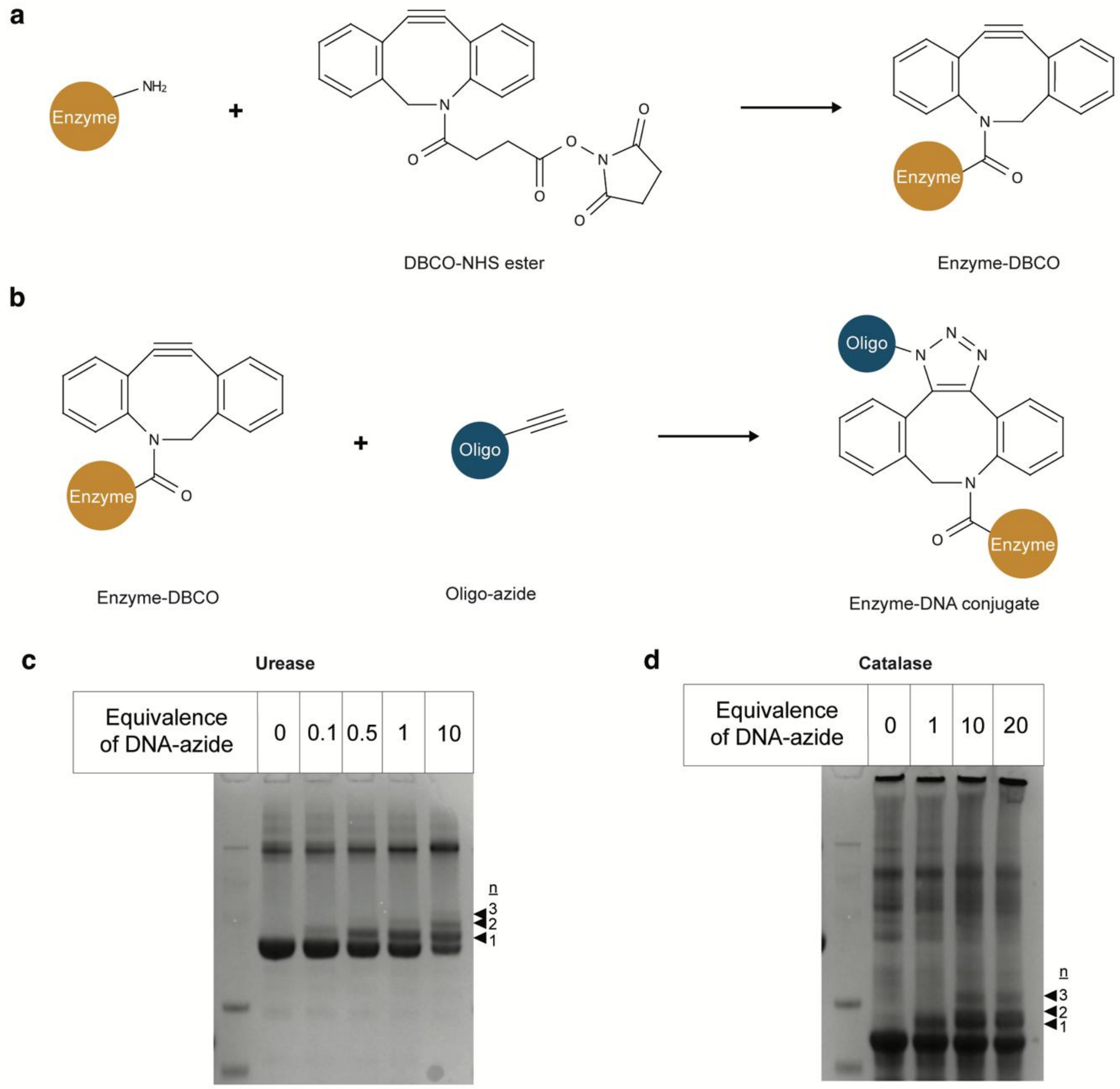

In this approach, the enzyme is first functionalized with a DBCO moiety via NHS esters coupling to accessible lysine residues. As the mono-labeled proteins are not purified prior to the subsequent DNA conjugation step, the presence of multiple DBCO groups can result in a heterogeneous product containing variable numbers of attached DNA strands. To balance DNA consumption with the yield of mono-labeled enzyme conjugates, a 1-fold molar excess of DNA was used for urease and a 5-fold excess for catalase. Following conjugation, the products were purified and concentrated using Amicon filtration and subsequently used for binding to the DNA origami structures.

**Supplementary Figure 8. BSA-flat sheet origami conjugates.**

(a) Schematic representation of flat sheet origami with five (Asym) or ten (Sym) protein capture strands, which were incubated overnight with varying molar equivalents of BSA-DNA conjugates. (b) Asym samples analyzed by 1% agarose gel electrophoresis (1x TBE, 12.5 mM MgCl2) showed comparable mobility shifts at 8- and 16-fold molar excess. Bands were excised and purified via Amicon filtration to remove unbound protein before AFM imaging. (c) Representative AFM image of samples with 16-fold excess; schematic indicates BSA positions (dots). Scale bar: 200 nm.(d) Sym samples analyzed similarly with 4- and 8-fold excess showing comparable shifts; bands excised and purified prior to AFM imaging; (e) Representative AFM image of samples with 8-fold excess; schematic shows BSA positions. Scale bar: 200 nm.

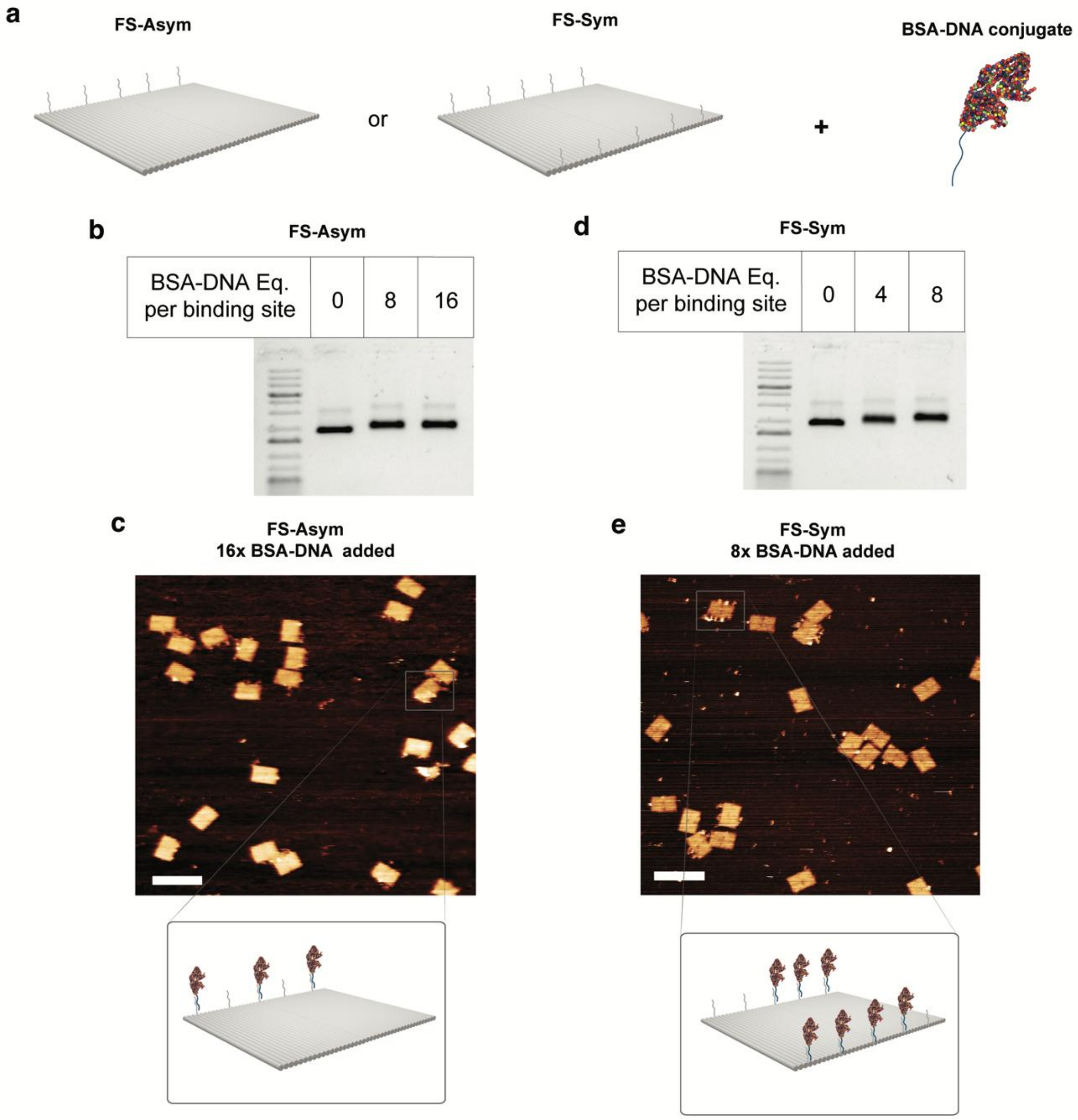

**Supplementary Figure 9. Urease-flat sheet origami conjugates.**

(a) Schematic representation of flat sheet origami with five (Asym) or ten (Sym) protein capture strands, which were incubated overnight with varying molar equivalents of urease-DNA conjugates. (b) Asym samples analyzed by 1% agarose gel electrophoresis (1x TBE, 12.5 mM $MgCl_2$) showed similar mobility shifts at 8-, 16-, and 40-fold excess; bands from the 8-fold sample were excised from the lower band (c) and upper band (d). (c,d) Full AFM images of the excised 8-fold sample bands, with a schematic indicating urease protein positions (dots). Scale bar: 200 nm. (e) Sym samples analyzed similarly; band from 20-fold excess excised. (f) AFM image of the 20-fold Sym sample excised from the lower band. A crop-out of both AFM images are used in Figure 3.

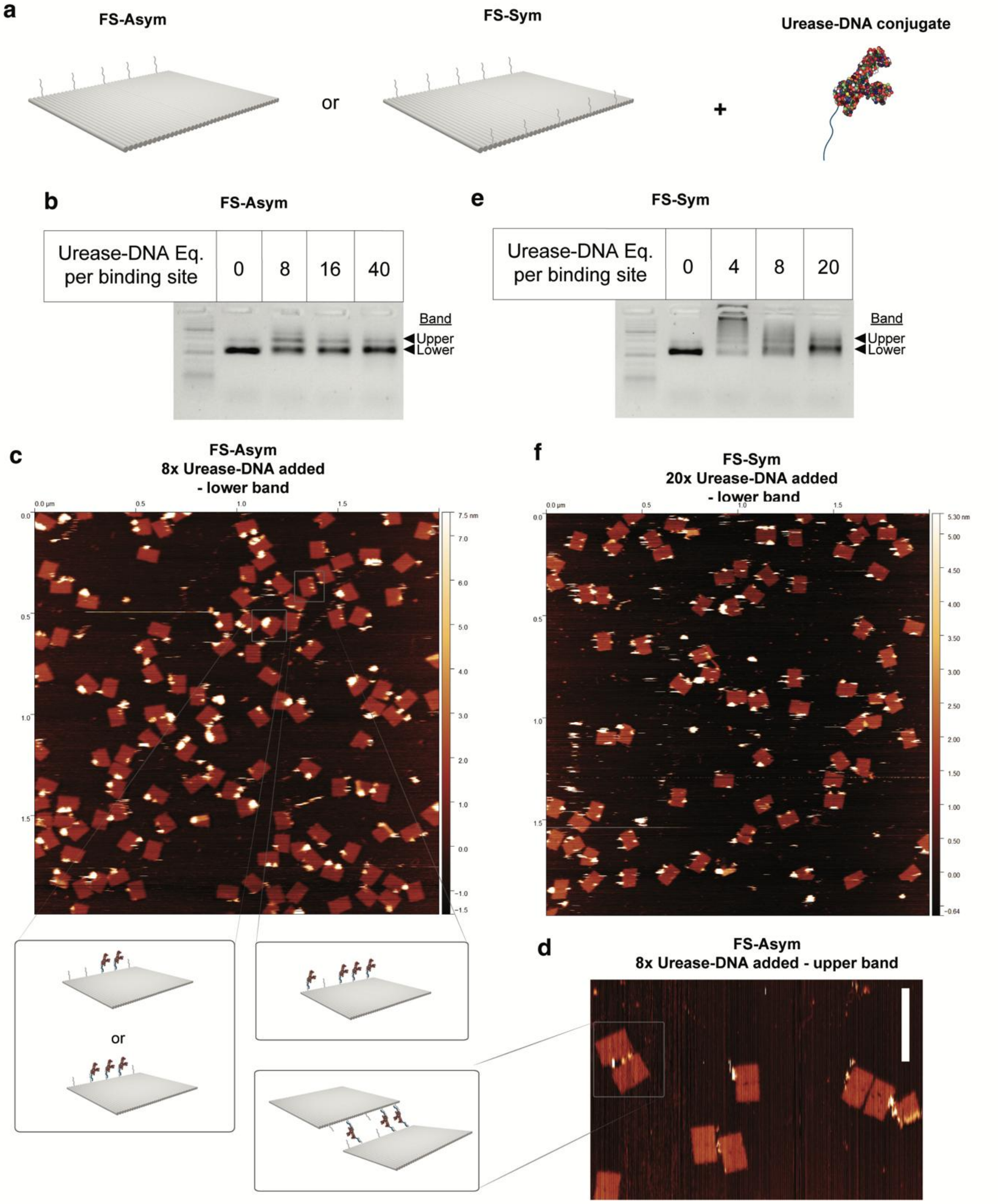

**Supplementary Figure 10. Catalase-flat sheet origami conjugates.**

(a) Schematic of flat sheet origami with five (Asym) or ten (Sym) protein capture strands incubated overnight with varying molar equivalents of catalase-DNA conjugates. (b) Samples analyzed by 1% agarose gel electrophoresis (1x TBE, 12.5 mM $MgCl_2$) showed comparable mobility shifts. Bands from the Sym design (ten strands) with 20-fold excess were excised. (c) Representative AFM image of the excised Sym sample. Scale bar: 200 nm. (d) Full AFM image of the excised Asym (8x equivalence) after a second gel purification step. A crop out of this image is used in Figure 4.

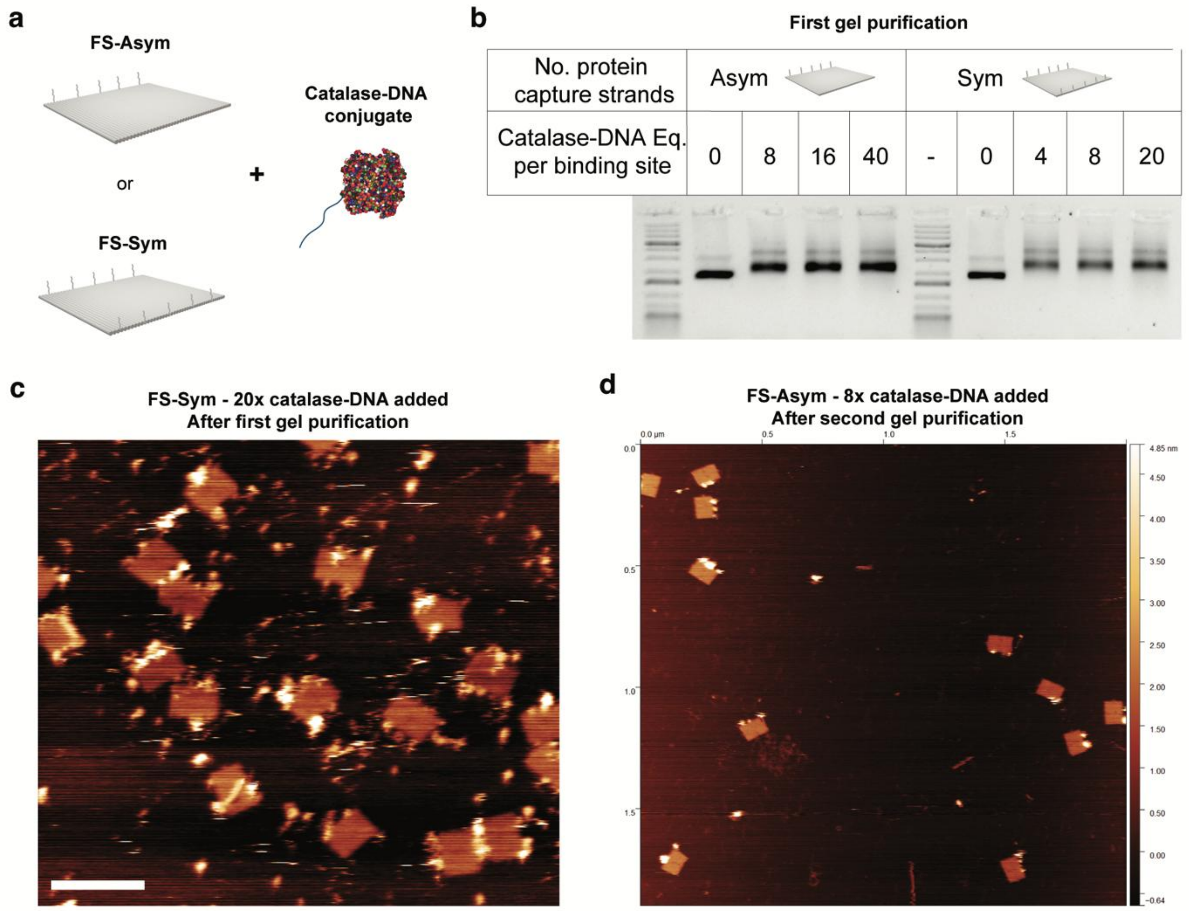

**Supplementary Figure 11. Color-coded diffusion tracks .**

It shows color-coded diffusion tracks from of Asym-catalase flat sheet origami at 100 mM $H_2O_2$, showing significant sample drift that compromises the reliability of particle tracking.

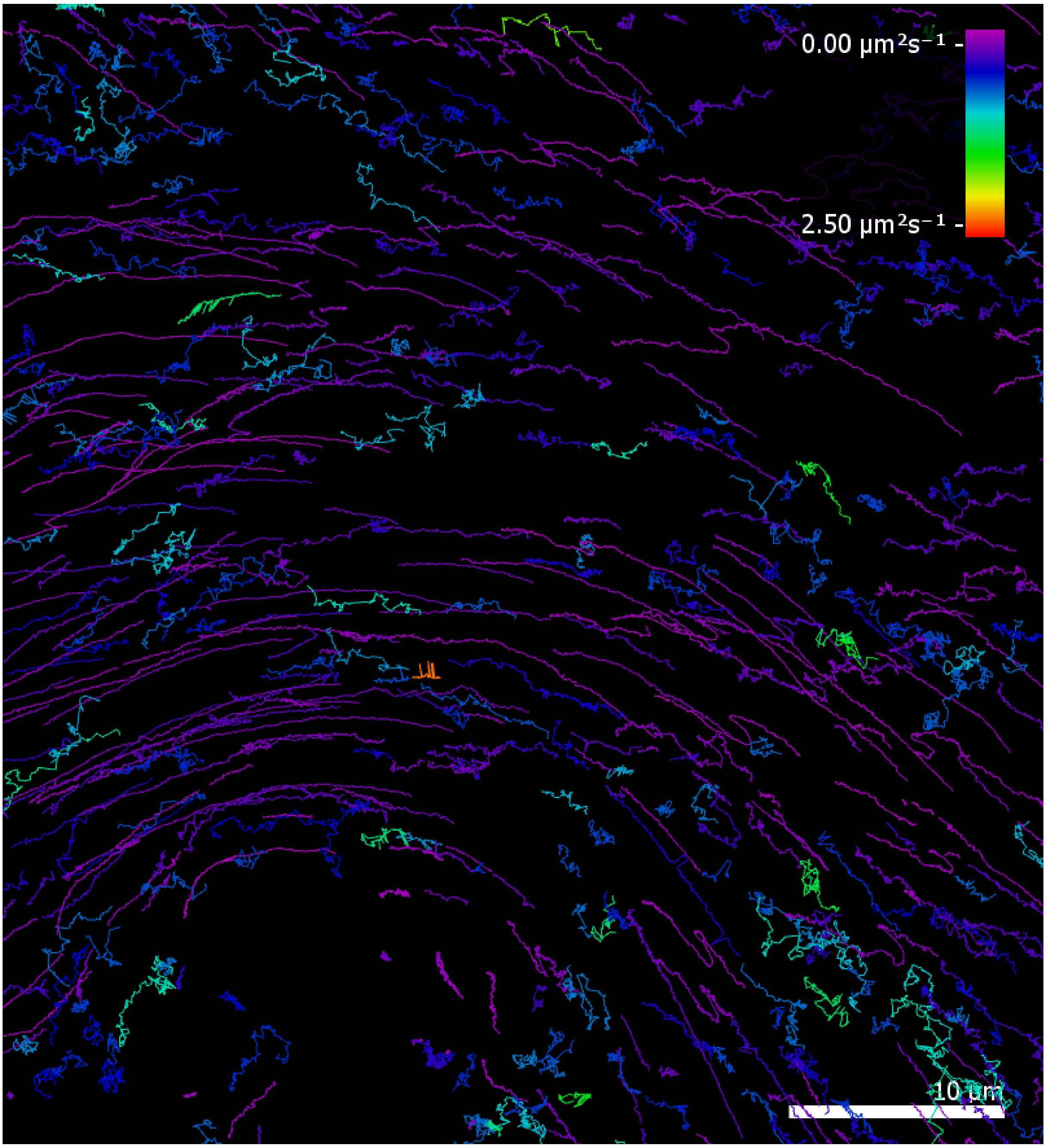

**Table 1. Table of core sequences for constructing the flat sheet DNA origami.**

| Start | Sequence |
|---|---|
| 13[48] | TGGATAGCTTACCAGACGACGATAAAACGAAC |
| 28[95] | AAGTGTATTTCACCAGTGAGACCCTGAGAG |
| 9[112] | TTGCCCTGAACATAAAAACAGGGACCTTTACA |
| 17[176] | AAAACTTTCCGGAATCATAATTACTTTAACAA |
| 28[63] | GCTAACTCGTTTGCGTATTGGGCGGGTTTGCC |
| 14[191] | AATAAACATGTAATTTAGGCAGAGTAAATAAG |
| 29[48] | GAGAGGCGACATTAATTGCGTTGCTCCCCGGG |
| 18[159] | TGAGAGAAGTACATAAATCAATAGATGATG |
| 5[176] | CTTGAGCCTCAGTAGCGACAGAATATCACCGG |
| 3[112] | AAAATCTCTCGGTCATAGCCCCCTGTTTTCAT |
| 5[208] | TCACCAGTCACCAATGAAACCATCCTCCCTCA |
| 3[48] | TCAGCGGACGTAACGATCTAAAGTTCAGAACC |
| 10[127] | GAGAGAATACGAGAAACACCAGAAAATAAGGC |
| 6[191] | AATCAATAATGTTAGCAAACGTAGTAGCAATA |
| 7[176] | TACGCAGTGAAAATTCATATGGTTGTCACCGA |
| 19[48] | GCCTTTATCAAAATTAAGCAATAACCATTAGA |
| 9[48] | ACAAGAACCGAACTGACCAACTTTTTTGACCC |
| 12[191] | GCCCAATATCATTCCAAGAACGGGACGACGAC |
| 24[63] | TCGTAACCCAGGCAAAGCGCCATTATTAAGTT |
| 9[144] | AGACGGGATAGCCGAACAAAGTTAGGAATACC |
| 10[31] | AGAACTGAAATCTACGTTAATAAAAACCAA |
| 18[191] | AGTCAATACCTTGCTTCTGTAAATGAATTATT |
| 14[159] | GCCTGTTTTGAGAATCGCCATATAGAAAAA |
| 20[159] | AAACAAATCAGGTTTAACGTCACTGAATAA |
| 17[208] | ATCTTCTGGTGATAAATAAGGCGTGCATTTTC |
| 29[80] | GTTTTTCTAAGCCTGGGGTGCCTAGGTCATAG |
| 1[112] | GTAACACTAATAAATCCTCATTAATTGATATT |
| 9[176] | AAGTCAGAACCGAAGCCCTTTTTACTCCTTAT |
| 19[176] | GTGAATAAGTGAATTTATCAAAATAACGCGAG |
| 28[127] | TTCACCAGCACAACATACGAGCCGTCCGCTCA |
| 5[144] | TTCATTAACGTCAGACTGTAGCGCTATTAGCG |
| 6[31] | CGAAGGCACTAAAACACTCATCGAAAGAGG |
| 3[80] | AGGAATTGTGTAGCATTCCACAGAATTTTCAG |
| 10[159] | AACGTCAATCAAGATTAGTTGCTAGAAGGC |
| 7[208] | TACATAAACACGGAATAAGTTTATCAGCAAAA |
| 9[80] | GTAACAAAAACGAGGCGCAGACGGCAAAGTAC |
| 28[31] | CCCGCTTTAATGAATCGGCCAATGGTGGTT |
| 29[112] | GCTGATTGAAAGAGTCTGTCCATCGTGAGGCC |
| 19[208] | TTAATTAAAGCTTAGATTAAGACGTTAATTTC |

| | |
|---|---|
| 10[63] | TTAATCATCAACATTATTACAGGTCTATCATA |
| 24[127] | TACAAACAAACAAACGGCGGATTGTCGGATTC |
| 20[191] | CATTTCAACAGTACCTTTTACATCTCAATATA |
| 6[159] | CCAAAGACTGGCATGATTAAGAAGAAAAGT |
| 18[31] | AGCATAACCTGTAATACTTTTGGATAAATT |
| 12[159] | TTATCCGAAACCAATCAATAATCAGAACGC |
| 22[159] | TGGAAGGACGTTATTAATTTTAAATAGATA |
| 10[191] | TCCCAATCACAATTTTATCCTGAAATTACCGC |
| 28[159] | TCGTCTGGTAGCAATACTTCTTGCCAGAAT |
| 29[144] | TAACCGTTAAATGGATTATTTACAAAAAGGGA |
| 10[95] | AAATTGGTGAGATTTAGGAATAAAAGGAAT |
| 7[48] | CCAGCGATCGGGTAAAATACGTAATCGCTGAG |
| 25[208] | CAGTTGGCATCTAAAATATCTTTAAAAGAAAC |
| 26[95] | AAAGGGGTGTGTGAAATTGTTAGAAGCATA |
| 27[80] | CTGTTTCCGATGTGCTGCAAGGCGCGCCATTC |
| 30[127] | ACCGAGTACCCTTCACCGCCTGGCGGGCAACA |
| 11[176] | ACCCAGCTCAAATAAGAAACGATTCCTGAACA |
| 23[48] | TGGCCTTCAATATTTTGTTAAAATGCCTGAGA |
| 16[191] | AATAAACATTCAAATATATTTTAGCTGAGAAG |
| 18[63] | AGAATTAGTTCAACGCAAGGATAACACCATCA |
| 2[159] | TGAGGCACTTTTCATAATCAAACAAGTTTG |
| 3[144] | TTTGCCATGGTCAGACGATTGGCCAGCCAGAA |
| 4[31] | TAGTTGCAACCGATATATTCGGTGCCACTA |
| 22[191] | ATCCTGATTATCATTTTGCGGAACGGAGCACT |
| 15[112] | TCATTTTTGTATAAAGCCAACGCTATACAAAT |
| 8[159] | AAGCAGAGAATTAACTGAACACTTTTGTTT |
| 11[80] | TCATCAGTGCTTGAGATGGTTTAAAAATCAAC |
| 4[127] | CGGCATTTCAAAAAAAAGGCTCCATCACGTTG |
| 16[95] | TGAATATTCATTTGGGGCGCGAAGTAGCAT |
| 24[159] | ATACATTTCTAAAGCATCACCTAGATAAAA |
| 25[144] | ATGAAAAATGAGGATTTAGAAGTATAAATCCT |
| 17[80] | CTATATTTAATGCTGTAGCTCAACGATTAGAG |
| 2[63] | ATAGTTAGGTGAGAATAGAAAGGACTTGCTTT |
| 23[80] | AAATGTGATTGTATAAGCAAATATTGAACGGT |
| 8[63] | AAGGGAACCGGATATTCATTACCCTTTCAACT |
| 5[48] | GCTTGCAGATTTCTTAAACAGCTTCAACAGTT |
| 26[127] | TGCAACAGTCTTCGCTATTACGCCGCGATCGG |
| 13[176] | TTTCCTTAGCAAGCAAATCAGATATATTTTGC |
| 16[127] | TCTTACCAGCGGATGGCTTAGAGCATAAGAGG |
| 23[176] | AGTAACATTGTTTGGATTATACTTGATGAATA |
| 25[48] | CCGGAAACGTGCATCTGCCAGTTTTTCGCGTC |
| 28[191] | GCTCATGGCACTTGCCTGAGTAGACCGATTAA |

| | |
|---|---|
| 29[176] | AATAACATAAATACCTACATTTTGCCCTTCTG |
| 4[159] | CCTTTAGAGGTGAATTATCACCTACCAGCG |
| 7[80] | AACGGAGAAAAGACTTTTTCATGATTTGCGGG |
| 15[144] | GGGCTTAATATCAACAATAGATAAATTTACGA |
| 16[31] | TCATTCCGTAGATTTAGTTTGAAGCCTCAG |
| 27[112] | CAATTCCATCACACGACCAGTAATTTGGCAGA |
| 11[208] | ACGCTAACCAAAATAAACAGCCATATATCAGA |
| 2[95] | CAACGCCCGAATAATAATTTTTAAAGGAGC |
| 18[95] | TAACATCCATATATTTTAAATGAAAGGGTG |
| 2[191] | CACCAGAGCCACCACCGGAACCGCGATAGCAG |
| 3[176] | AACCAGAGCCGCCGCCAGCATTGAACCGTTCC |
| 15[48] | CGGAAGCAAAAGCGGATTGCATCAGTTTAGAC |
| 23[208] | CACCAGAAAGATGATGGCAATTCAGGGAGAAA |
| 8[191] | GCTATCTTGGGTAATTGAGCGCTAATTATTTA |
| 21[48] | GTCTGGAGTTCAACCGTTCTAGCTCGGGAGAA |
| 7[112] | GTGTCGAAGAAACGCAATAATAACCCAGAAGG |
| 24[191] | AACAACTATCAAACCCTCAATCAACCATTAAA |
| 25[176] | CTCAAATAATAGATTAGAGCCGTCAAAGTTTG |
| 17[112] | GGTGGCATATATAACTATATGTAACGGCTTAG |
| 26[63] | GGGTAACGTCGAATTCGTAATCATATGAGTGA |
| 12[31] | AATAGCGGGGTAATAGTAAAATAAAAGATT |
| 22[31] | AATTTTTCGCCATCAAAAATAAGAGGGGAC |
| 23[112] | TCCGTGGGATTCGACAACTCGTATTTAGACTT |
| 16[159] | GCCTGTTAATCGCAAGACAAAGCATAGGTC |
| 8[95] | TAGCCGGGCTGCTCATTCAGTGCGAGTAGT |
| 27[144] | CATTCTGGGAGGCGGTCAGTATTAAGCAGCAA |
| 2[127] | CACAAACAGAGTTTCGTCACCAGTCATGTACC |
| 13[208] | CAAGTACCTTTCATCGTAGGAATCTCTTACCA |
| 25[112] | TGCGGGCCTGCCACGCTGAGAGCCACACCGCC |
| 25[80] | AGGCTGCGGTCACGTTGGTGTAGATCAACATT |
| 18[127] | GTTGGGTTCAATTCTACTAATAGTGCTGAAAA |
| 29[208] | AACTATCGGCCAGCCATTGCAACACACAGACA |
| 11[48] | TAACGGAATGTGAATTACCTTATGTAATCTTG |
| 24[95] | GGGATAGCAACTGTTGGGAAGGAGCTGGCG |
| 4[191] | CACCGTAAATTTGGGAATTAGAGCTTTGTCAC |
| 15[176] | CGCCAACAACATGTTCAGCTAATGCGGCTGTC |
| 16[63] | AATATGCAGCAAATGGTCAATAACCAAGGCAA |
| 17[48] | TACATTTCACTAAAGTACGGTGTCAACCAGAC |
| 15[80] | AGTACCTTAGGTCTTTACCCTGACTCGTCATA |
| 21[176] | TACAGTAATTACCTGAGCAAAAGAATATGTGA |
| 26[159] | CAGAGGTCCAACAGAGATAGAAACGCTCAA |
| 7[144] | CAAAAGAACAAAAGGGCGACATTCGGAAATTA |

| | |
|---|---|
| 27[48] | TACCGAGCCCAGGGTTTTCCCAGTTTCTGGTG |
| 26[191] | AATACCGAGCGTAAGAATACGTGGGGAAAAAC |
| 17[144] | GCAAATCCTAGTATCATATGCGTTCAACAGTA |
| 23[144] | TTGCCCGAGTTAGAACCTACCATAGAAATTGC |
| 5[80] | ATCGTCACTGTATCGGTTTATCAGACAACTAA |
| 24[31] | GACGACAGCTTTCCGGCACCGCCACGACGT |
| 27[176] | ACCTGAAAACGAACCACCAGCAGATGCTGAAC |
| 8[127] | AAACCGAGATCCGCGACCTGCTCCGATAAATT |
| 0[191] | TTAATGCCTCATACATGGCTTTTGAGAACCAC |
| 1[80] | GGATAGCACAGTACCAGGCGGATAAGTGCCGT |
| 30[63] | CCAGCAGGGAACAAGAGTCCACTATTAAAGAA |
| 1[208] | GGAGTGTAGAGTAACAGTGCCCGTATAAACAG |
| 31[48] | CCAGTTTGCGAAAATCCTGTTTGACGCGCGGG |
| 31[80] | CGTGGACTGCAAGCGGTCCACGCTCCAGGGTG |
| 1[176] | AGTAAGCGCCCTGCCTATTTCGGAACCTATTA |
| 30[95] | AGTTGCACCAACGTCAAAGGGCGAAAAACC |
| 31[112] | GTCTATCAGGCAAGTGTAGCGGTCACGCTGCG |
| 30[31] | CCGAAATCCGAGATAGGGTTGAGTGTTGTT |
| 1[144] | TGGAAAGCACATGAAAGTATTAAGAGGCTGAG |
| 31[144] | CGTAACCAAGTGTTTTTATAATCAACGCAAAT |
| 1[48] | GCCACCCTTTGATATAAGTATAGCCCGGAATA |
| 30[159] | CCTGAGACCACACCCGCCGCGCTTAATGCG |
| 0[31] | GGTGTATTCAGAACCGCCACCCTTTGTCGT |
| 31[176] | CCGCTACATAGACAGGAACGGTACTGATTAGT |
| 0[63] | CGAGAGGGCAGAGCCACCACCCTCCAGCCCTC |
| 31[208] | CGAGCACGGGAGCTAAACAGGAGGAGAACTCA |
| 0[95] | TTTTGCTAGCCCAATAGGAACCACAAACTA |
| 30[191] | AGGGATTTGGGCGCGTACTATGGTTGCTTTGA |
| 0[127] | ACTCCTCAAGAGAAGGATTAGGATTAGCGGGG |

**Table 2. Table of staple sequences for the attachment of Cy3 or ATTO647N to the flat sheet DNA origami.**

For the FRET experiment, 15 staples with an extension for capturing the Cy3-fuctionalized DNA strand was used. For the single molecule tracking experiments, 6 staples with an extension for capturing the ATTO647N-functionalized DNA strand was used. In this case, staples 1-6 with the extension (in red) and staples 7-15 without the capture extension (thus, is unfunctionalized -UF) were used for annealing the structure.

| Name | Sequence |
|---|---|
| UF Cy3/ATTO 1 | CCATTAAATATACCAAGCGCGAAATCAATCAT |
| UF Cy3/ATTO 2 | TCAGAAGCAACTCCAACAGGTCAGATGTTTTA |
| UF Cy3/ATTO 3 | TAAACGTTCTGTAGCCAGCTTTCATGGGCGCA |
| UF Cy3/ATTO 4 | AATATTCACATAGTAAGAGCAACAAGAAAGAT |
| UF Cy3/ATTO 5 | AATCGTAACCGGAGACAGTCAAATAAATTTTT |
| UF Cy3/ATTO 6 | GAGGACTTTTGTATCATCGCCTATGTTACT |
| UF Cy3 7 | AAAAATCTAATTGCTCCTTTTGTTAATTGC |
| UF Cy3 8 | AGGAAGAGCGAGTAACAACCCGACCGTAAT |
| UF Cy3 9 | TAGCGAACCAGATACATAACGCCACCACATTC |
| UF Cy3 10 | ATTTAACATGTAGGTAAAGATTCACAATGCCT |
| UF Cy3 11 | GAGGGAGGGGTAGCAACGGCTACAACAGCATC |
| UF Cy3 12 | CAAGAAAAAGAAAACGAGAATGACATGCTTTA |
| UF Cy3 13 | ATTTGCACGATAATCAGAAAAGCCCATATGTA |
| UF Cy3 14 | GCATGTAGGTATTCTAAGAACGCGGGTTTTGA |
| UF Cy3 15 | GTAGATTTCATCAAGAAAACAAAACTTTTTTA |
| Cy3/ATTO 1 | CCATTAAATATACCAAGCGCGAAATCAATCATGGGCTCATGCGAGGCTGTATGT |
| Cy3/ATTO 2 | TCAGAAGCAACTCCAACAGGTCAGATGTTTTAGGGCTCATGCGAGGCTGTATGT |
| Cy3/ATTO 3 | TAAACGTTCTGTAGCCAGCTTTCATGGGCGCAGGGCTCATGCGAGGCTGTATGT |
| Cy3/ATTO 4 | AATATTCACATAGTAAGAGCAACAAGAAAGATGGGCTCATGCGAGGCTGTATGT |
| Cy3/ATTO 5 | AATCGTAACCGGAGACAGTCAAATAAATTTTTGGGCTCATGCGAGGCTGTATGT |
| Cy3/ATTO 6 | GAGGACTTTTGTATCATCGCCTATGTTACTGGGCTCATGCGAGGCTGTATGT |
| Cy3 7 | AAAAATCTAATTGCTCCTTTTGTTAATTGCGGGCTCATGCGAGGCTGTATGT |
| Cy3 8 | AGGAAGAGCGAGTAACAACCCGACCGTAATGGGCTCATGCGAGGCTGTATGT |
| Cy3 9 | TAGCGAACCAGATACATAACGCCACCACATTCGGGCTCATGCGAGGCTGTATG T |
| Cy3 10 | ATTTAACATGTAGGTAAAGATTCACAATGCCTGGGCTCATGCGAGGCTGTATGT |
| Cy3 11 | GAGGGAGGGGTAGCAACGGCTACAACAGCATCGGGCTCATGCGAGGCTGTAT GT |
| Cy3 12 | CAAGAAAAAGAAAACGAGAATGACATGCTTTAGGGCTCATGCGAGGCTGTATGT |
| Cy3 13 | ATTTGCACGATAATCAGAAAAGCCCATATGTAGGGCTCATGCGAGGCTGTATGT |
| Cy3 14 | GCATGTAGGTATTCTAAGAACGCGGGTTTTGAGGGCTCATGCGAGGCTGTATGT |
| Cy3 15 | GTAGATTTCATCAAGAAAACAAAACTTTTTTAGGGCTCATGCGAGGCTGTATGT |
| Cy3 strand | ACATACAGCCTCGCATGAGCCC-/3AmMC6T/ |

| | |
|---|---|
| ATTO647N strand | ACATACAGCCTCGCATGAGCCC-/3ATTO647NN/ |

**Table 3. Table of staple sequences for the attachment of Cy5 to the flat sheet DNA origami.**

For the FRET experiment, 15 staples with an extension for capturing the Cy5-fuctionalized DNA strand was used. In this case, all staples 15 with the extension (in blue) were used for annealing the structure. For the single molecule tracking experiments, where only an ATTO647N was needed, the 15 unfunctionalized staples (UF), which lacked the capture strand), were used.

| Name | Sequence |
|---|---|
| UF Cy5 1 | CGAGGTGAGGAGTTAAAGGCCGCTGGAAGTTT |
| UF Cy5 2 | ACCCTCGTGTCCAATACTGCGGAATATTATAG |
| UF Cy5 3 | ATATGATACAAACAAGAGAATCGATTAAATTG |
| UF Cy5 4 | TCATCAGTGCTTGAGATGGTTTAAAAATCAAC |
| UF Cy5 5 | AGAACCCTCAATAAATCATACAGGCTGTTTAG |
| UF Cy5 6 | CTTTAATCCTCAGCAGCGAAAGGAGGCTTT |
| UF Cy5 7 | TACGAGGTTGAATCCCCCTCAACATAAATC |
| UF Cy5 8 | AGAAAGGAACTAGCATGTCAATCCAAAAAC |
| UF Cy5 9 | AACTAATGCTCCCGACTTGCGGGAAGGCGTTT |
| UF Cy5 10 | GAGTAATGATTTCATTTGAATTACTTAATTAC |
| UF Cy5 11 | GGAACGAGGAAGGTAAATATTGACAACCGATT |
| UF Cy5 12 | AACAGTTCATAATATCCCATCCTAGTCCTGAA |
| UF Cy5 13 | CCCCGGTTGTAAAACAGAAATAAATCAAAATT |
| UF Cy5 14 | AGCCTTAAAAAATGAAAATAGCAGAGCGCATT |
| UF Cy5 15 | ATGGAAACCTACCTTTTTAACCTCATGCTGAT |
| Cy5 1 | TGGCTGTGGTGTGCGAGTAACTCGAGGTGAGGAGTTAAAGGCCGCTGGAAGTTT |
| Cy5 2 | TGGCTGTGGTGTGCGAGTAACTACCCTCGTGTCCAATACTGCGGAATATTATAG |
| Cy5 3 | TGGCTGTGGTGTGCGAGTAACTATATGATACAAACAAGAGAATCGATTAAATTG |
| Cy5 4 | TGGCTGTGGTGTGCGAGTAACTTCATCAGTGCTTGAGATGGTTTAAAAATCAAC |
| Cy5 5 | TGGCTGTGGTGTGCGAGTAACTAGAACCCTCAATAAATCATACAGGCTGTTTAG |
| Cy5 6 | TGGCTGTGGTGTGCGAGTAACTCTTTAATCCTCAGCAGCGAAAGGAGGCTTT |
| Cy5 7 | TGGCTGTGGTGTGCGAGTAACTTACGAGGTTGAATCCCCCTCAACATAAATC |
| Cy5 8 | TGGCTGTGGTGTGCGAGTAACTAGAAAGGAACTAGCATGTCAATCCAAAAAC |
| Cy5 9 | TGGCTGTGGTGTGCGAGTAACTAACTAATGCTCCCGACTTGCGGGAAGGCGTTT |
| Cy5 10 | TGGCTGTGGTGTGCGAGTAACTGAGTAATGATTTCATTTGAATTACTTAATTAC |
| Cy5 11 | TGGCTGTGGTGTGCGAGTAACTGGAACGAGGAAGGTAAATATTGACAACCGATT |
| Cy5 12 | TGGCTGTGGTGTGCGAGTAACTAACAGTTCATAATATCCCATCCTAGTCCTGAA |
| Cy5 13 | TGGCTGTGGTGTGCGAGTAACTCCCCGGTTGTAAAACAGAAATAAATCAAAATT |
| Cy5 14 | TGGCTGTGGTGTGCGAGTAACTAGCCTTAAAAAATGAAAATAGCAGAGCGCATT |
| Cy5 15 | TGGCTGTGGTGTGCGAGTAACTATGGAAACCTACCTTTTTAACCTCATGCTGAT |
| Cy5 strand | AGTTAC/iCy5/TCGCACACCACAGCCA |

**Table 4. Table of staple sequences for the attachment of protein-DNA conjugates to the flat sheet DNA origami.**

When no proteins are needed on the flat sheet, all 10 unfunctionalized (UF) staples were used. For structures with five capture strands (FS-Asym configuration), staples 1-5 with the capture extension (in orange) were used, along with unfunctionalized (UF) staples 6-10. For structures with ten capture strands (FS-Sym configuration), staples 1-10 with capture extensions were used.

| Name | Sequence |
|---|---|
| UF Enzyme 1 | CTTTCCATTGCTAAACAACTTTGATACCGA |
| UF Enzyme 2 | ACAGATGGACCTTCATCAAGAGCGATTTTA |
| UF Enzyme 3 | AAGAGGATCGAGCTTCAAAGCGTGGAAGTT |
| UF Enzyme 4 | AATGCCGGGCTATCAGGTCATTTCGCATTA |
| UF Enzyme 5 | TGTAAAAGTCGACTCTAGAGGAGCTCACTG |
| UF Enzyme 6 | GAGCCGCCCCTCAGAGCCGCCACCATGATACA |
| UF Enzyme 7 | GAGATAACGCAAGAAACAATGAAAAAAATACA |
| UF Enzyme 8 | GAGCCAGTAAGTAATTCTGTCCAGTATTAAAC |
| UF Enzyme 9 | CAATAACGAAAATCGCGCAGAGGCCGTCGCTA |
| UF Enzyme 10 | ATATTTTTTAGCCCTAAAACATCGTATCTGGT |
| Enzyme 1 | CTTTCCATTGCTAAACAACTTTGATACCGAAAAATTCAGGATTCTCAATT |
| Enzyme 2 | ACAGATGGACCTTCATCAAGAGCGATTTTAAAAATTCAGGATTCTCAATT |
| Enzyme 3 | AAGAGGATCGAGCTTCAAAGCGTGGAAGTTAAAATTCAGGATTCTCAATT |
| Enzyme 4 | AATGCCGGGCTATCAGGTCATTTCGCATTAAAAATTCAGGATTCTCAATT |
| Enzyme 5 | TGTAAAAGTCGACTCTAGAGGAGCTCACTGAAAATTCAGGATTCTCAATT |
| Enzyme 6 | GAGCCGCCCCTCAGAGCCGCCACCATGATACAAAAATTCAGGATTCTCAATT |
| Enzyme 7 | GAGATAACGCAAGAAACAATGAAAAAAATACAAAAATTCAGGATTCTCAATT |
| Enzyme 8 | GAGCCAGTAAGTAATTCTGTCCAGTATTAAACAAAATTCAGGATTCTCAATT |
| Enzyme 9 | CAATAACGAAAATCGCGCAGAGGCCGTCGCTAAAAATTCAGGATTCTCAATT |
| Enzyme 10 | ATATTTTTTAGCCCTAAAACATCGTATCTGGTAAAATTCAGGATTCTCAATT |
| Protein strand - NHS | /5AmMC6/ AATTGAGAATCCTGAATTTT |
| Protein strand - Azide | /5AzideN/ AATTGAGAATCCTGAATTTT |

**Table 5. Table of core sequences for constructing the 14HB DNA origami.**

| Start | Sequence |
|---|---|
| 0[83] | CCAGTACCAACTTTGGTTTATCAGCTTGGCCGCTTATCCCTTTTATTTC |
| 0[104] | CCGTAACGAGTGAGAAAGGAGCCTTTAACCCTCAGGGTGGTTAATTTTT |
| 0[146] | CCCTCATTTTCAGGGTAAAGCACTAAATGCAAGCG |
| 0[167] | CCGCCACTTGCCTTAAACCATCGATAGCAAAGACTCTGAGAGAAGGGTG |
| 0[209] | GGAGGTTTCGGTCAGTAGCACCATTACCACGTAATCAGTGAGGATATTC |
| 0[258] | GAGAGGGTTGATATTCAAAATCCGACTT |
| 0[272] | GGATAAGCCACCACAAAGGTGAATTATCTCATCTTATCGGCCAAAGTGT |
| 0[335] | TTAAGAGCCCTCAGACAAAAGGGCGACACGCCTGACATGGAATTTGACG |
| 0[398] | CCCGTATAGGTCAGCCACGGAATAAGTT |
| 0[440] | TGGTAATTCATTAAACATACATAAAGGTCGCAATAATTAGTAGACCTGA |
| 0[461] | TTTGATGGCGCAGTATGTTAGCAAACGTAAAAGAACGTTGTAGTGGCAC |
| 1[28] | GTCTTTCTAGCGTATAAAGAACGTGGACGTTGAGTTATATTCATTATTA |
| 0[83] | CCAGTACCAACTTTGGTTTATCAGCTTGGCCGCTTATCCCTTTTATTTC |
| 0[104] | CCGTAACGAGTGAGAAAGGAGCCTTTAACCCTCAGGGTGGTTAATTTTT |
| 0[146] | CCCTCATTTTCAGGGTAAAGCACTAAATGCAAGCG |
| 0[167] | CCGCCACTTGCCTTAAACCATCGATAGCAAAGACTCTGAGAGAAGGGTG |
| 0[209] | GGAGGTTTCGGTCAGTAGCACCATTACCACGTAATCAGTGAGGATATTC |
| 0[258] | GAGAGGGTTGATATTCAAAATCCGACTT |
| 0[272] | GGATAAGCCACCACAAAGGTGAATTATCTCATCTTATCGGCCAAAGTGT |
| 0[335] | TTAAGAGCCCTCAGACAAAAGGGCGACACGCCTGACATGGAATTTGACG |
| 0[398] | CCCGTATAGGTCAGCCACGGAATAAGTT |
| 0[440] | TGGTAATTCATTAAACATACATAAAGGTCGCAATAATTAGTAGACCTGA |
| 0[461] | TTTGATGGCGCAGTATGTTAGCAAACGTAAAAGAACGTTGTAGTGGCAC |
| 1[28] | GTCTTTCTAGCGTATAAAGAACGTGGACGTTGAGTTATATTCATTATTA |
| 10[314] | TTGCTGATTTATCCAGAAGGC |
| 11[35] | ATTATGACAATAAACTAATGCCGAGCTTCAAAGCGGATTAGAGCTGTAG |
| 11[42] | CCCTGTAAGATAGGTCCAACGACAGCCCTCATAGTCAGACGTGATACCG |
| 11[56] | TGCGGGACAAAGAAAAAAGGATTCAAAT |
| 11[77] | AACGCAACAATAAATAAGAGCTTAAGAGGAAGCCC |
| 11[98] | AGAACCCAATAGTACCTCGTTAAGCGGA |
| 11[119] | TGCAATGAAAGGTGAAAAACCTGACTATTATAGTCAAATTTAGTAAATG |
| 11[126] | CCTGAGTCTGGTTTAGGTGCCGATAGCAAGCCCAAACTAAAGATCTCCA |
| 11[140] | AGGTAAAGATTCAAAGTTGCACGGAACCGCCACCA |
| 11[161] | AGAAAGGGGATTTACAGAGGGGAAAACG |
| 11[203] | AACCGTTAACAACTATACTGCTAAATAT |
| 11[245] | CGAGCCGAAGGAATCCAACGCGTTTTGAAGCCTTATAAAGCCTCCTTGA |
| 11[266] | AAAGCCTAGTTGGCCCAGAGCGAACCTCCCGACTTGGGCTTATTAATTA |
| 11[308] | GCGTTGCGCTACATATACCTCACTGCCCGAGATTTCAGGCGC |
| 11[329] | CTCAATCTGAAAAAAACGATTAATAGCA |
| 11[350] | ATTTACAACGCTGAAAAAATGGTAGGAATCATTACAAAGTACTCATTTG |
| 11[371] | AGTCACAAACACCGTACAGAGCAAGCCGTTTTTATGTAATTCAATTAAT |

| | |
|---|---|
| 11[392] | AGGGACAAGAGGTGACAGGGAACCGCAC |
| 11[434] | AAGCGTAGCCATTAAACAAAGCTGTCTT |
| 11[455] | AGACAATAACTGATAGCGCTATAGAAACCAATCAAACAAGAAAAGTTAC |
| 12[83] | CGGCAAATTGCGGGGGAAGAAAAATCTAGGCATAGTCATACATTCCCAA |
| 12[237] | TGCGTATGGAGCGGGCCCGGAATAGGTGGCGTTTGTTAGAGC |
| 12[384] | TTGCTGGGGAGCTATAATGCCCCCTGCCGCATTGATCACAAT |
| 13[56] | GCGAAAAAAAAGAAGCAGGGAAAAACGA |
| 13[98] | ATCACCCGTTTGATCAGCGAAATTATAC |
| 13[119] | GGGGTCGGCCCCAGCGAGGGTTGCGATT |
| 13[161] | GAGCCCCCCTGGCCTTTTCATGAGATGGTTTAATTGTTTTGC |
| 13[182] | ACGGGGAACAGCTGTAAACGGAACGAGT |
| 13[203] | GGCGAGATTTTCACGCCACTATGCCCTG |
| 13[224] | AGCGAAATGGGCGCTAAAACGGCTCATTCAGTGAA |
| 13[245] | GGCGCTGGGGGAGAATACACTCCAAATC |
| 13[266] | CACGCTGTTAATGATGACCCCCAAGAAC |
| 13[287] | ACCCGCCAAACCTGAGCGCGACTTCATC |
| 13[308] | GCTACAGGGCGCGTCTCCTCAAGAGAAGCCGCCACGATTGAG |
| 13[329] | TTGCTTTAAACGCTTAAATTGACAGATG |
| 13[350] | ACGTGCTCGCCAGCACCTGCTACTGACC |
| 13[357] | TTCCTCGGAACCTATTATTCTCCACCACTATGGTT |
| 13[371] | CAGAGCGTAATATCCGGAACGGGTCAAT |
| 13[392] | GGCCGATTCAAACTAGCCGAACCCTTTT |
| 13[413] | ACAGGAACACTTGCGGAAACCGCAATAG |
| 13[434] | CCTGAGATTCTTTGATAACGGAGAGCAA |
| 13[455] | CAGTGAGAATTAACCTGGCATATTGAGT |
| 2[55] | TTCTTAAGAGGCTTTAGCCCGATACTTT |
| 2[118] | AAAAAAAATCGGAACAGGCGATTTTAAA |
| 2[181] | AAACGTCTTTCCATATTGCCCCAGTCAA |
| 2[286] | GGAAATTTATACCATCGTGCCCTAATGA |
| 2[349] | TACCAGCATCCGCGCATTGCAATGGATT |
| 2[412] | GAAACGCACCAGAACTGAGTACAACAGA |
| 11[182] | CTTTAAAGAATCATAAATCAT |
| 11[189] | CGAGGCGTATTTAATGCTTCT |
| 11[287] | GTATTAAAATGCAGCCTGAGC |
| 11[294] | TCAATATACGGGCAAAGCCGG |
| 11[413] | AACTCACAGTCGGGGCGCTTA |
| 11[420] | CCCTTCTATAACATCGGTACG |

**Table 6. Table of staple sequences for the attachment of ATTO647N to the 14HB DNA origami.**

For the single molecule tracking experiments, 6 staples with an extension for capturing the ATTO647N-functionalized DNA strand was used. In this case, staples 1-6 with the extension (in red) were used for annealing the structure. If no ATTO fluorophores need to be captured, the unfunctionalized staples (UF) with no capture extension were used.

| Start | Sequence |
|---|---|
| UF ATTO 1 | ACTAACGTAACGCCTTAGCAATTCATTC |
| UF ATTO 2 | TTAAGAACGACGATGCATCAATACATTT |
| UF ATTO 3 | AGTAAATGTAAAATGATAATATAAATCC |
| UF ATTO 4 | AAGAGTAGCCAGTTATCAATAATTATCA |
| UF ATTO 5 | AACTTTGTAACGTCGAGCCAGGGAAGGG |
| UF ATTO 6 | CTATCTTTAGACGGGCAGAAGTAGATTT |
| ATTO 1 | ACTAACGTAACGCCTTAGCAATTCATTCGGGCTCATGCGAGGCTGTATGT |
| ATTO 2 | TTAAGAACGACGATGCATCAATACATTTGGGCTCATGCGAGGCTGTATGT |
| ATTO 3 | AGTAAATGTAAAATGATAATATAAATCCGGGCTCATGCGAGGCTGTATGT |
| ATTO 4 | AAGAGTAGCCAGTTATCAATAATTATCAGGGCTCATGCGAGGCTGTATGT |
| ATTO 5 | AACTTTGTAACGTCGAGCCAGGGAAGGGGGGCTCATGCGAGGCTGTATGT |
| ATTO 6 | CTATCTTTAGACGGGCAGAAGTAGATTTGGGCTCATGCGAGGCTGTATGT |
| ATTO647N strand | ACATACAGCCTCGCATGAGCCC-/3ATTO647NN/ |

**Table 7. Table of staple sequences for the attachment of protein-DNA conjugates to the flat sheet DNA origami.**

When no enzymes are needed on the flat sheet, all 10 unfunctionalized (UF) staples were used. For structures with five capture strands (14HB-Asym), staples 1-5 with the capture extension (in orange) were used, along with unfunctionalized (UF) staples 6-10. For structures with ten capture strands (14HB-Sym), staples 1-10 with capture extensions were used.

| Start | Sequence |
|---|---|
| UF Enzyme 1 | TAGTAAATGAATTTTCCACAGTCAAAGG |
| UF Enzyme 2 | AATAGAAAGGAACATAGGAACGTTTTTT |
| UF Enzyme 3 | TAGCGTCAGACTGTCGCCACCGAGCTTG |
| UF Enzyme 4 | CGGAACCGCCTCCCGCTCAGTCCACCAC |
| UF Enzyme 5 | AGCCGCCACCAGAAGAAACATACGTATA |
| UF Enzyme 6 | ACGATTGGCCTTGACTTGAGTATTTTAG |
| UF Enzyme 7 | GCTAAAAAACTACAACGCCTGTAGCATTCTGTATGGTGAAT |
| UF Enzyme 8 | ATCACCAGTCAATAGTTTAGATCAAATG |
| UF Enzyme 9 | ATTAGCGGGGTTTTTCAGAGCTATTGAC |
| UF Enzyme 10 | AACGGGGTCAGTGCTATTCACTATAAAA |
| Enzyme 1 | TAGTAAATGAATTTTCCACAGTCAAAGGAAAATTCAGGATTCTCAATT |
| Enzyme 2 | AATAGAAAGGAACATAGGAACGTTTTTTAAAATTCAGGATTCTCAATT |
| Enzyme 3 | TAGCGTCAGACTGTCGCCACCGAGCTTGAAAATTCAGGATTCTCAATT |
| Enzyme 4 | CGGAACCGCCTCCCGCTCAGTCCACCACAAAATTCAGGATTCTCAATT |
| Enzyme 5 | AGCCGCCACCAGAAGAAACATACGTATAAAAATTCAGGATTCTCAATT |
| Enzyme 6 | ACGATTGGCCTTGACTTGAGTATTTTAGAAAATTCAGGATTCTCAATT |
| Enzyme 7 | GCTAAAAAACTACAACGCCTGTAGCATTCTGTATGGTGAATAAAATTCAGGATTCTCAATT |
| Enzyme 8 | ATCACCAGTCAATAGTTTAGATCAAATGAAAATTCAGGATTCTCAATT |
| Enzyme 9 | ATTAGCGGGGTTTTTCAGAGCTATTGACAAAATTCAGGATTCTCAATT |
| Enzyme 10 | AACGGGGTCAGTGCTATTCACTATAAAAAAAATTCAGGATTCTCAATT |
| Protein strand - NHS | /5AmMC6/ AATTGAGAATCCTGAATTTT |
| Protein strand - Azide | /5AzideN/ AATTGAGAATCCTGAATTTT |
